# Dysbiosis of a leaf microbiome is caused by enzyme secretion of opportunistic *Xanthomonas* strains

**DOI:** 10.1101/2023.05.09.539948

**Authors:** Sebastian Pfeilmeier, Anja Werz, Marine Ote, Miriam Bortfeld-Miller, Pascal Kirner, Andreas Keppler, Lucas Hemmerle, Christoph G. Gäbelein, Christine M. Pestalozzi, Julia A. Vorholt

## Abstract

Dysbiosis is characterized by a perturbed microbiota associated with host disease. In both plants and animals, the innate immune system contributes to maintain microbiota homeostasis in healthy organisms, with NADPH oxidases playing a crucial role. In *Arabidopsis thaliana*, the absence of NADPH oxidase RBOHD can lead to an altered leaf microbiota, including an enrichment of opportunistic *Xanthomonas* pathogens. It is currently unclear whether the microbiota change occurs independently of the opportunistic pathogens or is caused by the latter, and which virulence factors of *Xanthomonas* are essential for its opportunistic lifestyle. Here, we found that the opportunistic *Xanthomonas* strains secrete a cocktail of cell wall degrading enzymes via the type-2 secretion system (T2SS) that degrade leaf tissue and promote *Xanthomonas* growth during plant infection. Both disease severity and leaf degradation activity were increased in *rbohD* compared to Col-0 plants, attesting to the opportunistic behaviour of the *Xanthomonas* strains on immune compromised plants. Using gnotobiotic plant experiments with a synthetic bacterial community of more than 100 commensal strains and drop-in of *Xanthomonas* wildtype or mutant strains revealed that T2SS-dependent virulence is required for plant disease and for the shift in microbiota composition. Overall, our data indicate that a single opportunistic pathogen can drive community shifts, here caused by tissue damage in leaves, creating an environment in which specific commensal bacteria can thrive.

## Introduction

Host-associated microbial communities, collectively referred to as microbiota, promote development, growth, and adaptation to abiotic and biotic stress in healthy host organisms. Bacteria are highly abundant members in the microbiota and assemble into taxonomically structured communities in animals and plants^1-3^

Under certain circumstances, the relationship between host and its microbiota can become unbalanced, resulting in an alternative state of the microbial community termed dysbiosis, which is commonly associated with disease and with an alteration in the composition or function of the microbiome^4,5^. The host immune system plays a central role in maintaining and controlling microbiota homeostasis to prevent dysbiosis^4,6^. In addition, opportunistic pathogens are particularly relevant in dysbiosis as they are normally harmless for the host but are equipped with potential virulence functions and, under conducive conditions, eventually cause context-dependent diseases. For example, the human opportunistic pathogens *Helicobacter pylori, Clostridium difficile* and *Staphylococcus epidermidis* show contextual pathogenicity in the gastrointestinal tract and on the skin, respectively, and are common members of the microbiota remaining asymptomatic in most cases^7,8^. Therefore, dysbiosis has underlying contributions both from individual species with pathogenic potential as well as from the microbiota.

Dysbiosis has also been described in the plant leaf microbiota^9,10^. A reverse genetic screen in *Arabidopsis thaliana* mutants with defects in the immune system revealed that *rbohD* knockout plants, among others, harbour an altered phyllosphere microbiota and develop disease^9^. In this case, two *Xanthomonas* strains, named Leaf131 and Leaf148, were identified as opportunistic pathogens in *rbohD* plants and as the driver of plant disease after inoculation with a bacterial synthetic community (SynCom) of an initial 222 strain community^9^ that was assembled from the At-LSPHERE strain collection^1^. Both *Xanthomonas* strains lack a type-3 secretion system, a typical virulence factor of *bona fide* pathogens, which might render them non-virulent in *A. thaliana* Col-0 wildtype, in the context of a microbiota.

In plants, the NADPH oxidase RBOHD produces apoplastic reactive oxygen species (ROS) and is involved in several pathways related to growth, development, and stress response^11-13^. Moreover, RBOHD is an important component of the plant immune system^14^. Plants recognize microbial activity due to the release of microbe- or danger-associated molecular patterns (MAMP or DAMP) or microbial effector proteins that lead to activation of RBOHD, which is a convergence point of pattern-triggered immunity (PTI) and effector-triggered immunity (ETI) signalling pathways^15^. RBOHD-produced ROS also function in cell wall polymer crosslinking during pathogen-induced lignification^16,17^. Apart from plants, other multi-cellular organisms possess NADPH oxidases, including fungi, where they serve both defence and differentiation signalling^18^, and mammals^11,19^, where they are involved in gut epithelial immune responses and prevent intestinal dysbiosis^20,21^.

To understand the mechanisms of how the microbiota is linked to plant health and disease, establishing causality is an essential premise. SynCom experiments have emerged as a decisive tool to study the processes and interactions shaping the microbiota and affecting the host^22^. For example, removal and late-introduction of individual strains have revealed priority effects due to the arrival order of strains in the phyllosphere microbiota of *A. thaliana*^23^. In another study, interactions between bacteria in a SynCom were mapped *in planta* and the underlying molecular mechanism of one of them characterized^24^. Using the SynCom approach, it has been demonstrated that certain taxa of the microbiota can have drastic impacts on host health and development^9,25,26^.

In this study, we dissect the contribution of opportunistic *Xanthomonas* strains, their context-dependent virulence, and host genotype to the bacterial community composition in the phyllosphere of *A. thaliana* using a SynCom approach and both targeted and random bacterial mutagenesis. We identified one of the two T2SS, Xps, as a major virulence factor of the opportunistic *Xanthomonas* Leaf131 and Leaf148 due to the secretion of plant tissue degrading enzymes. Furthermore, SynCom experiments revealed that *rbohD* does not directly affect the bacterial community but causes the shift in the microbiota of *rbohD* plants indirectly via *Xanthomonas* Leaf131 activity. Our results highlight the importance of the T2SS for opportunistic *Xanthomonas* pathogens during plant colonization as it affects plant health and the composition of the commensal microbiota.

## Results

### Disease and microbiota shift caused by the opportunistic *Xanthomonas* Leaf 131 on *rbohD* plants

*A. thaliana* plants with defective RBOHD, but not wild type plants, show impaired growth when inoculated with a synthetic community and exhibit a dysbiotic microbiota. The RBOHD phenotype can be remediated by removing the *Xanthomonas* Leaf131 strain from a 137-member microbiota community^9^. To determine whether the opportunistic pathogen not only drives plant disease but also alters the microbiota composition in *rbohD* plants, we inoculated germ-free *A. thaliana* seedlings with a SynCom of 137 strains that did or did not include *Xanthomonas* Leaf131 and analysed the community composition on Col-0 wildtype, *rbohD* knockout, and the complementation line *rbohD/RBOHD* by 16S rRNA amplicon sequencing.

As an indicator for monitoring overall community changes, we used effect size to quantify how much of the total variance in the microbiota is explained by the test factor, here plant genotype, and the significance of the effect was measured using PERMANOVA. As expected, we observed that the microbiota composition in *rbohD* plants compared to Col-0 was significantly altered when *Xanthomonas* Leaf131 was included in the microbiota, e.g. SynCom-137+Leaf131 with an effect size of 12.5% (p=0.0001). In contrast, the community composition did not significantly change when Leaf131 was omitted from the SynCom-137 (effect size 2.8%, p=0.72) (Figure 1A). Consistent with this, the difference in community composition of SynCom-137 in *rbohD* plants was observed when *Xanthomonas* Leaf131 was included but not in absence of the opportunistic pathogen, as indicated by a principal component analysis (Figure S1A). By analysing the changes in relative abundance of each strain in the SynCom-137, we found that specific strains were enriched in *rbohD* compared to Col-0, resulting in the characteristic microbiota shift in diseased *rbohD* plants as observed previously^9^. In addition to *Xanthomonas* Leaf131, we found the Gammaproteobacteria *Pseudomonas* Leaf58, Leaf127 and Leaf434, the Alphaproteobacterium *Sphingobium* Leaf26, the *Bacteroides* Pedobacter Leaf41, as well as the Actinobacterium *Sanguibacter* Leaf3 being enriched in their relative abundance (Figure 1B and Figure S1C). Here, we show that none of the changes in the relative abundance of these strains could be observed in *rbohD* plants in the absence of *Xanthomonas* Leaf131, which is also a Gammaproteobacterium.

**Figure 1.**
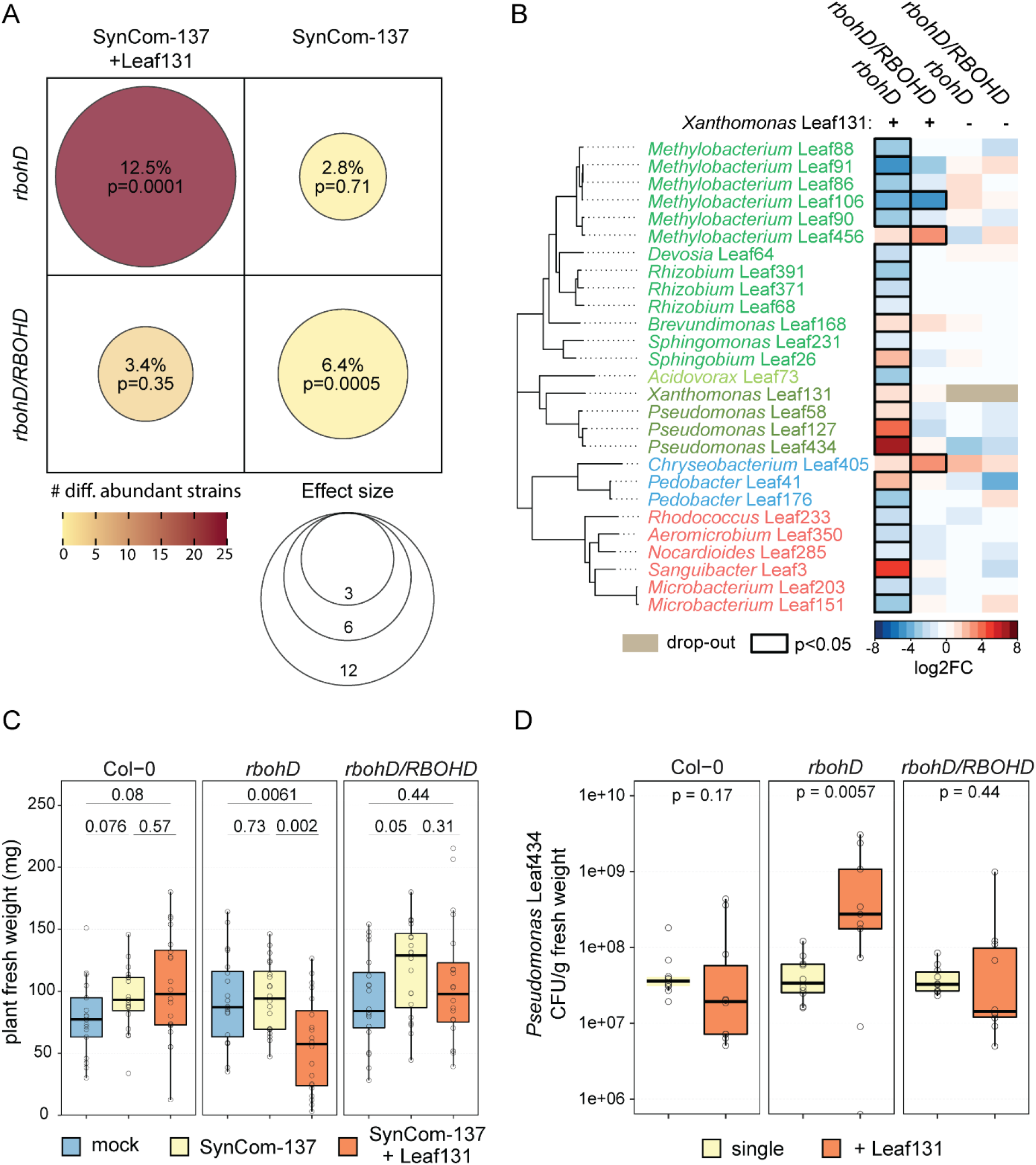
Microbiota shift and plant disease is driven by *Xanthomonas* Leaf131 in *rbohD* knockout plants. **A)** Composition of synthetic bacterial communities SynCom-137+ *Xanthomonas* Leaf131 or SynCom-137 in *rbohD* or *rbohD*/*RBOHD* plants was compared to Col-0 wild-type plants. Effect size represents percentage of total variance explained by genotype (shown by dot size and absolute value) and statistical significance is expressed with p-values determined by PERMANOVA (Benjamini– Hochberg adjusted, n = 16). Number of differentially abundant strains (as shown in panel B) is shown by dot colour. **B)** Heatmap shows subset of strains in SynCom-137 with significant log_2_ fold changes (log2FC, p-value < 0.05) in *rbohD* or *rbohD*/*RBOHD* compared to Col-0 wild-type plants in the presence (+) or absence (-) of *Xanthomonas* Leaf131. Black rectangles show significant changes, p-value < 0.05 (n = 16, Wald test, Benjamini–Hochberg adjusted). Complete heatmap of all strains in SynCom-137 is shown in Figure S1C. **C)** Fresh weight of aboveground plant tissue of Col-0, *rbohD* and *rbohD/RBOHD* mock inoculated or with SynCom-137 or SynCom-137+*Xanthomonas* Leaf131. Box plots show the median with upper and lower quartiles and whiskers present 1.5× interquartile range (n = 16, two-tailed Mann–Whitney *U*-test, p-values indicated above box plots). Corresponding plant phenotypes shown in Figure S1D. **D)** Colony forming units (CFU) counts of *Pseudomonas* Leaf434 per gram plant fresh weight after inoculation of germ-free Col-0, *rbohD* and *rbohD/RBOHD* plants with *Pseudomonas* Leaf434 as single inoculation or in binary inoculation with *Xanthomonas* Leaf131. Box plots show the median with upper and lower quartiles and whiskers present 1.5× interquartile range (n = 12, two-tailed Mann–Whitney *U*-test, p-values indicated above box plots).

As the SynCom-137 did not show significant differences in community composition in *rbohD* compared to the control Col-0 without *Xanthomonas* Leaf131 (Figure 1A and B and Figure S1A and B), we conclude that *rbohD* does not affect the microbiota *per se*, but rather indirectly via *Xanthomonas* Leaf131. Consistently, only *rbohD* knockout plants showed disease symptoms and a reduced average plant fresh weight after inoculation with SynCom137+Leaf131, but not Col-0 or *rbohD/RBOHD* (Figure 1C and Figure S1D).

To exemplarily validate that certain member of the microbiota benefit from the presence of *Xanthomonas* Leaf131 on *rbohD* plants but not on Col-0 wild type plants, we selected a commensal strain, *Pseudomonas* Leaf434, that was enriched on *rbohD* plants based on our data from the SynCom experiment (Figure 1B) and assessed its absolute abundance in a binary inoculation experiment together with *Xanthomonas* Leaf131. Substantiating the results of the SynCom experiment, *Pseudomonas* Leaf434 showed higher plant colonization levels only in *rbohD* plants when inoculated together with *Xanthomonas* Leaf131 compared to single inoculation or in control Col-0 and *rbohD/RBOHD* plants (Figure 1D).

Overall, our data show that the presence of the conditional pathogen *Xanthomonas* Leaf131 promotes the growth of specific microbiota members in *rbohD* plants.

### Plant tissue degradation by *Xanthomonas* Leaf131 and Leaf148

When examining possible virulence mechanisms, we found that *Xanthomonas* Leaf131 and also Leaf148, previously identified as opportunistic pathogens^9^, degrade leaf tissue. We therefore set up a quantitative *A. thaliana* assay to assess tissue degradation using leaf discs (Figure 2). Both *Xanthomonas* strains degraded the tissue, resulting in translucent leaf discs, which we quantified using pixel brightness (Figure 2A and B). We observed that tissue degradation was markedly more severe in leaf discs of *rbohD* plants compared to Col-0 plants (Figure 2A and B), corroborating the stronger disease phenotype of these *Xanthomonas* strains in *rbohD* plants^9^.

**Figure 2.**
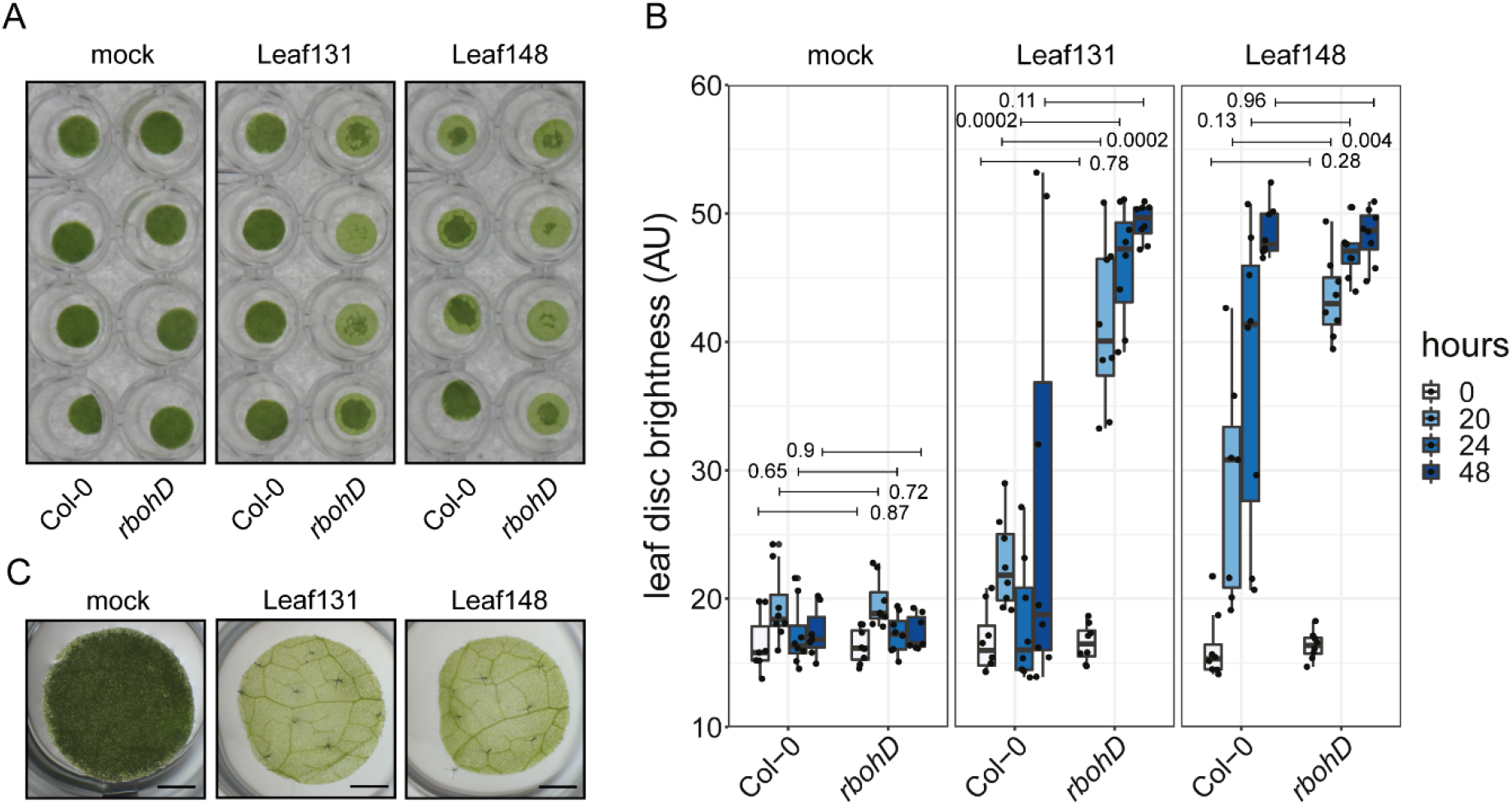
*Xanthomonas* Leaf131 and Leaf148 degrade plant tissue. **A)** Leaf discs of Col-0 and *rbohD* plants (six weeks old) were mock inoculated (10 mM MgCl_2_) or with *Xanthomonas* Leaf131 or Leaf148 (OD=0.02) and incubated for 20 hours. **B)** Time-course measurement and quantification of leaf disc brightness (arbitrary unit, AU) from experiment described in A). Statistical differences between Col-0 and *rbohD* at varying time points is indicated above box plots with p-value on right or left of horizontal line indicating comparison (two-tailed Mann–Whitney *U*-test, n = 8). Box plots show the median with upper and lower quartiles and whiskers present 1.5x interquartile range. **C)** Leaf disc of five-week-old *rbohD* plants mock (10 mM MgCl_2_) inoculated or with *Xanthomonas* Leaf131 or Leaf148 (OD=0.02) and incubated for 48 hours. Scale bar depicts 1 mm.

Leaf tissue degradation progressed gradually over time starting at the edges of the leaf discs (Figure 2B and S2A). After complete degradation of leaf tissue, the leaf disc was translucent and eventually lost its cellular cohesion and fragmented after mechanical impact (Figure S2B). In contrast to the effective degradation of leaf discs from *rbohD* plants, those from Col-0 plants showed reduced and patchy degradation even after 48 hours (Figure S2C). We tested other plant genotypes impaired in PTI signalling upstream of RBOHD and found that the triple co-receptor mutant *bak1/bkk1/cerk1* (*bbc*) but not the triple receptor mutant *fls2/efr/cerk1* (*fec*) was susceptible to leaf disc degradation similar to *rbohD* (Figure S3B). Beside plant genotype, plant age influenced leaf disc degradation with five-week-old Col-0 plants being more vulnerable compared to six-week-old plants (Figure S3A) suggesting that several plant factors affect the phenotype. While we found that intact leaves remained unaffected upon exposure to *Xanthomonas* Leaf131 and Leaf148 within two days of observation, wounded leaves showed visible signs of degradation over the same time period, as expected due to more accessible tissue (Figure S2D).

### Secretion of a cocktail of cell wall degrading enzymes via one of two T2SS by *Xanthomonas*

Leaf tissue degradation by *Xanthomonas* as a proxy for a virulence phenotype was observed not only by viable bacteria but also by cell-free supernatants of liquid cultures (Figure 3A), indicating that the phenotype is mediated by secreted factors. Consistent with this finding, the secretion of cell wall degrading enzymes (CWDE) by the T2SS is a known virulence function of other *Xanthomonas* species^27,28^. *Xanthomonas* Leaf131 and Leaf148 each possess two T2SS gene clusters, designated *xps* and *xcs* by homology search. To test whether the degradation activity is dependent on the T2SS, we deleted the core genes of the two T2SS operons (Figure 3B) and generated double mutants in *Xanthomonas* Leaf131 and Leaf148. In both strains, the *xps* mutant and the double knockout *xpsxcs* did not show tissue degradation, in contrast to the *xcs* mutants, which were still able to degrade leaf discs (Figure 3C and D). This indicates that the T2SS Xps is required for leaf degradation by *Xanthomonas*, which is in line with studies of other *Xanthomonas* bacteria reporting the importance of *xps* for virulence^29-31^.

**Figure 3.**
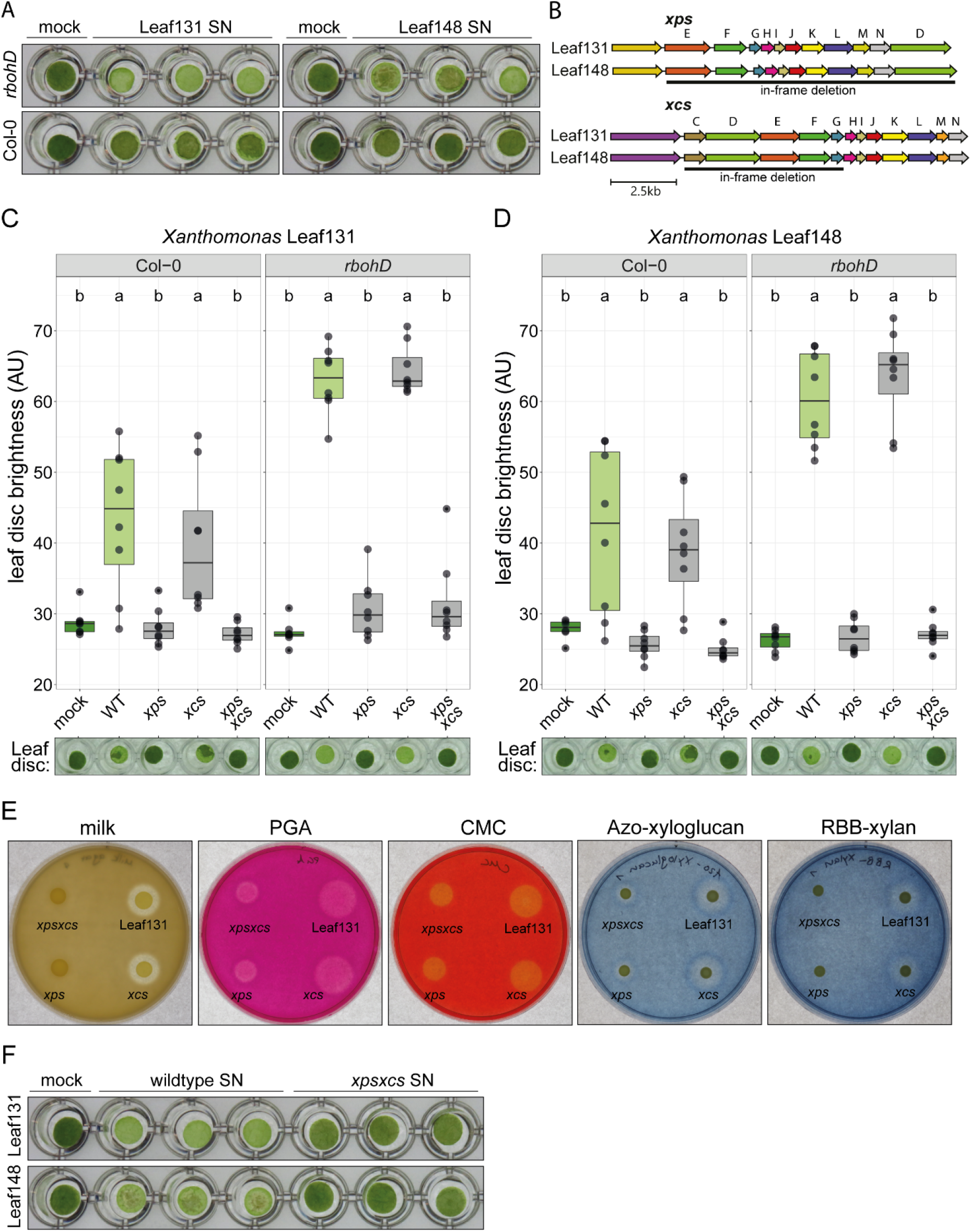
Type-2 secretion system Xps is required for leaf tissue degradation and secretion of plant polymer-degradative enzymes. **A)** Leaf discs of Col-0 and *rbohD* plants (five weeks old) were mock treated (0.5x LB) or treated with cell-free supernatant (0.22 μm filter-sterilized) of *Xanthomonas* Leaf131 or Leaf148 liquid cultures and incubated for 48 hours. **B)** Genomic region of the T2SS operons *xps* and *xcs* in *Xanthomonas* Leaf131 and Leaf148. Letters indicate gene names and black line shows region of gene deletion. **C)** Leaf discs of Col-0 or *rbohD* plants (six weeks old) were mock treated or with *Xanthomonas* wildtype or mutant strains of Leaf131 or **D)** Leaf148 and incubated for 24 hours. Significant differences were calculated with ANOVA and Tukey’s HSD post hoc test (n = 8, letters indicate significance groups, *α* = 0.05). **E)** Agar plates containing either skimmed milk, polygalacturonic acid (PGA), carboxymethyl cellulose (CMC), Azo-xyloglucan and Remazol Brilliant Blue-Xylan (RBB-Xylan). Drops of 4 μl *Xanthomonas* Leaf131 wildtype or mutant suspension were pipetted onto agar plate. Pictures were taken 24 hours after incubation at 22°C. **F)** Leaf discs of five-week-old *rbohD* plants were mock treated (0.5x LB) or treated with 0.22 μm filter-sterilized supernatant of liquid cultures from *Xanthomonas* Leaf131 or Leaf148 wildtype and *xpsxcs* mutants. Leaf discs were incubated for 48 hours at 22°C.

In addition, we deleted the *hrpX* and *hrpG* genes in *Xanthomonas* Leaf131, which encode regulators of various virulence factors including T2SS-secreted enzymes in various *Xanthomonas* pathogens^32-35^. However, the *hrpXhrpG* double knockout mutant still showed leaf degradation activity (Figure S4) suggesting that the production or secretion of the degradative enzymes is not, or not exclusively, controlled by HrpX or HrpG in *Xanthomonas* Leaf131.

To validate the finding that Xps is the primary T2SS involved in the secretion of plant polymer-degrading enzymes, we conducted agar plate assays using substrates for cell wall degrading enzymes (CWDE) with specific colour indicators. We tested *Xanthomonas* Leaf131 and found that the strain was able to degrade milk powder, polygalacturonic acid, carboxymethyl cellulose, xyloglucan and xylan, suggesting secretion of proteases, pectate lyases, glucanases and xylanases, respectively, as shown by halos that formed around the bacterial colonies after incubation indicating substrate degradation (Figure 3E). Notably, the T2SS mutants *xps* and *xpsxcs* showed reduced or delayed polymer degradation, unlike the *xcs* mutant strain (Figure 3E). However, *xps* and *xpsxcs* mutants still showed a small halo indicating substrate degradation, which might be due to cell lysis or alternative secretion mechanisms, such as outer membrane vesicles ^36^.

Next, we tested the leaf degradation activity of supernatants from *Xanthomonas* grown in liquid culture. As hypothesized, cell-free supernatant of *Xanthomonas* Leaf131 and Leaf148 T2SS mutant *xpsxcs* did not degrade *rbohD* leaf discs, in contrast to wildtype supernatant (Figure 3F). Analysing the total proteins of these *Xanthomonas* supernatants using SDS-PAGE, we noticed the absence of protein bands with a size between 35 to 55kDa in the *xpsxcs* supernatants that were present in supernatants from the wildtype strains (Figure S5A). We applied LC-MS/MS to identify the corresponding protein fractions that showed T2SS-dependent secretion (Supplementary table 1). Several candidate proteins are predicted to harbour a secretion signal peptide and a function potentially involved in plant interaction (Figure S5D). This included an endoglucanase (ASF73_13775), a serine protease (ASF73_18370), two pectate lyases (ASF73_04230, ASF73_20170) and a lysyl endopeptidase (ASF73_20190), which is in line with the activities observed in the agar plate assays. We generated in-frame deletion knockout strains in *Xanthomonas* Leaf131, and a gene cluster knockout including two genes located in proximity in the genome (Figure S5B) and tested the mutant strains for their leaf tissue degradation activity. Degradation was not affected in these mutant strains compared to wildtype in *rbohD* leaf discs (Figure S5C). Some of the mutant strains showed a difference in degradation in Col-0 (Figure S5C); however, this difference was not observed consistently, as leaf degradation in Col-0 is, in general, less pronounced, slower and more variable compared to *rbohD* (Figure 2B and D, Figure S2).

Overall, our data suggest that *Xanthomonas* secretes a cocktail of potential CWDE via the T2SS Xps that are responsible for leaf degradation.

### Identification of virulence factors of *Xanthomonas* using a forward genetic screen

In addition to using a targeted approach by mutating T2SS and genes for proteins that we found excreted under *in vitro* conditions, we used an untargeted approach by setting up a forward genetic screen in *Xanthomonas* Leaf131. We used a vanillate-inducible, hyperactive himar transposase (pAK415-himar)^37^ and tested more than 6000 transposon (Tn) mutants individually for their ability to degrade *rbohD* leaf discs. For all Tn mutants with delayed or impaired leaf degradation activity, we mapped the transposon insertion site by amplifying flanking regions using a nested-PCR approach with arbitrary PCR primers, followed by BLAST search with the sequenced PCR product^37^. After the primary screening, we validated 34 Tn mutant candidates on eight replicate leaf discs of *rbohD* to confirm their phenotypes and selected a subset of 16 Tn mutants based on their impairment in leaf disc degradation (Figure S6 and Supplementary table 2). Five Tn mutants lost leaf degrading capacity (Tn28_rplQ, Tn37_gtf, Tn68_dsbB, Tn80_xpsE, Tn81_xpsD) and five Tn mutants showed reduced leaf disc degradation (Tn10_glucanase, Tn15_iroN, Tn59_mopB, Tn69_flhA, Tn77_lolA) after 48 hours of co-incubation of bacteria and leaf discs, while six Tn mutants showed a delay of leaf disc degradation in *rbohD* only after 24 hours (Tn11_fliM, Tn13_tldD, Tn21_flgI, Tn24_dgkA, Tn26_XCC3185, Tn29_pilY) (Figure S6). The screening procedure was effective, as we identified Tn mutants which we had already confirmed as being important, e.g. the T2SS *xps*, and by identifying multiple independent transposon insertions in the same gene, suggesting high coverage (Supplementary table 2).

The Tn mutants with strong impairment of leaf degradation at 48 hours encoded genes involved in protein secretion (Tn80_xpsE, Tn81_xpsD), motility (Tn69_flhA), post-translational protein modification (Tn68_dsbB, Tn37_gtf), predicted glucanase/lectin domain (Tn10_glucanase), transport (Tn15_iroN) and protein translation (Tn28_rplQ). In summary, the screen resulted in several promising candidate genes of *Xanthomonas* involved in leaf degradation.

### T2SS Xps and additional virulence factors are required for full virulence during plant infection

To test the importance of the T2SS for virulence *in planta*, we inoculated Col-0 and *rbohD* plants with *Xanthomonas* Leaf131 wildtype and the T2SS mutants. Plant health was monitored by assessing disease symptoms and measuring plant fresh weight three weeks after inoculation using an established gnotobiotic growth system^9^. The infection experiment revealed that the virulence of *Xanthomonas* Leaf131 was dependent on the presence of the T2SS Xps, while Xcs did not contribute to virulence (Figure 4A and B), corroborating the results of the leaf degradation assay (Figure 3C). While the *Xanthomonas* Leaf131 *xps* mutant was non-virulent in Col-0 plants, as indicated by similar plant weight compared to mock inoculation, the *xps* mutant showed residual virulence in *rbohD* plants (Figure 4B), which suggests the presence of additional T2SS-independent virulence factors or alternative secretion pathways of leaf degrading enzymes. Moreover, the overall colonization level of these attenuated T2SS mutants *xps* and *xpsxcs* was reduced approximately one order of magnitude compared to *Xanthomonas* Leaf131 wildtype in Col-0 and *rbohD* plants (Figure 4C), highlighting the importance of the T2SS Xps for bacterial fitness during plant colonization.

**Figure 4.**
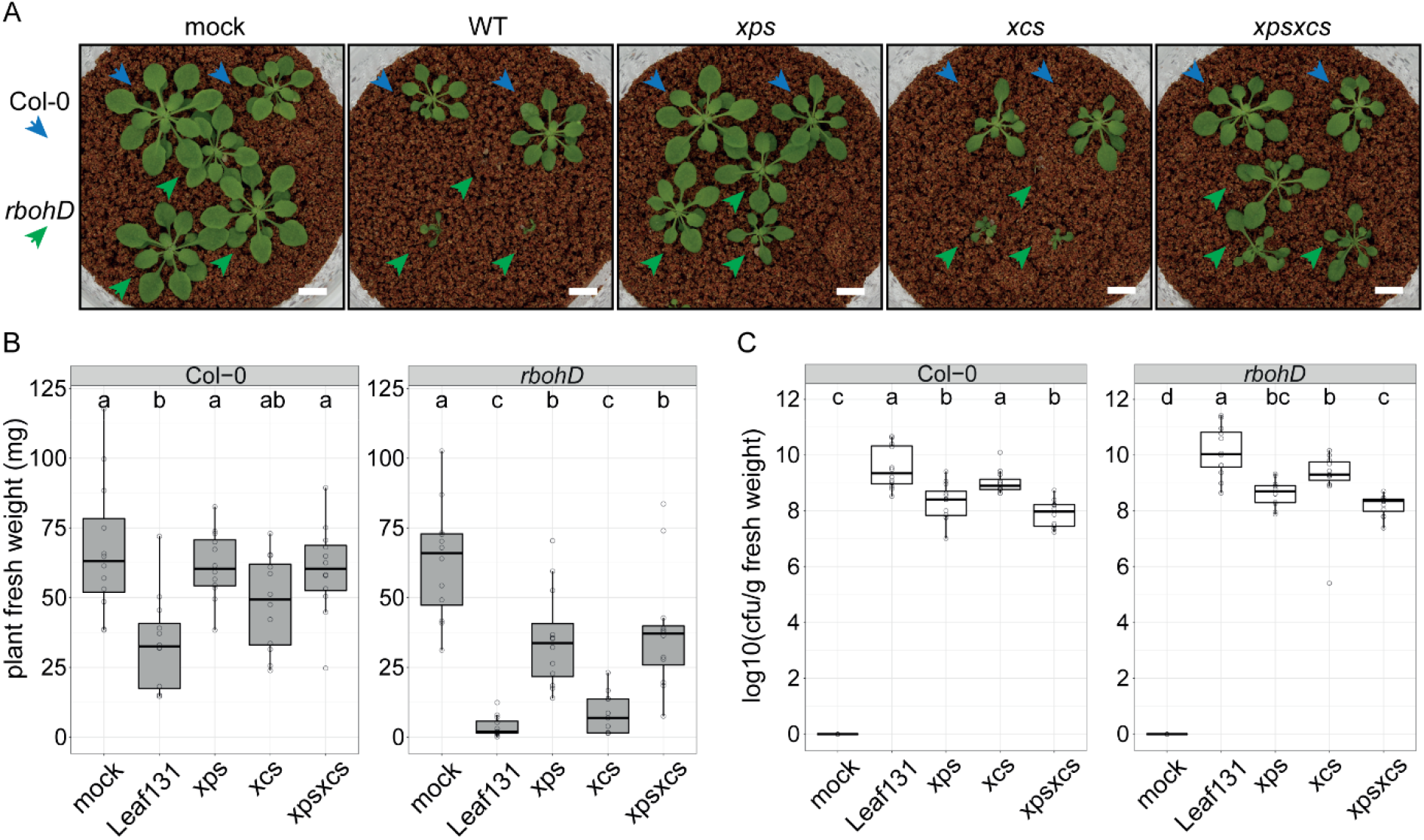
Only T2SS Xps is required for full virulence and fitness of *Xanthomonas* Leaf131 *in planta*. **A)** Phenotype of five-week-old Col-0 plants (blue arrow) and *rbohD* plants (green arrow) mock inoculated or with *Xanthomonas* Leaf131 wildtype (WT) or T2SS mutants *xps, xcs* and *xpsxcs*. **B)** Measurement of fresh weight from plants shown in A). **C)** Colony forming unit (CFU) counts of *Xanthomonas* Leaf131 per gram plant fresh weight from samples in B). Box plots show the median with upper and lower quartiles and whiskers present 1.5× interquartile range). Significant differences in B) (n=20) and C) (n=12) were calculated with ANOVA and Tukey’s HSD post hoc test (letters indicate significance groups, *α* = 0.05).

In addition, we tested the above identified Tn mutants for their virulence *in planta*. Seven out of 16 tested Tn mutants were less virulent than *Xanthomonas* Leaf131 wildtype during plant infection (Figure S7B). A number of Tn mutants with defects in motility related genes (e.g. Tn21_flgI, Tn29_pilY, Tn11_filM, Tn69_flhA) had no significant impact on *rbohD* plant weight compared to *Xanthomonas* Leaf131 wildtype despite a delay in leaf tissue degradation (Figure S6 and S7B). A potential explanation for this discrepancy could be that motility or adhesion to the leaf disc is critical for colonization or degradation activity up to 48 hours in liquid, but not relevant for virulence during plant colonization over a three-week-period. In addition, Tn mutants with reduced impact on plant weight also showed reduced absolute abundance on the host, as indicated by approx. 5-fold reduced cfu per gram fresh weight, similar as observed for the T2SS mutants (Figure S7C).

To validate the results of the Tn screen, we generated and tested mutants in candidate genes based on their phenotypes in leaf degradation and their virulence *in planta*. The selected targets were *dsbB* (ASF73_01480) encoding for the thiol-disulfide interchange protein; a glycosyltransferase (*gtf*, ASF73_08425), the encoding gene of which is located upstream of the flagellum gene cluster; a *glucanase* (ASF73_19940) encoding for a hypothetical protein with glucanase/lectin domain. In addition, we deleted a gene cluster including the operon encoding the identified glucanase, a TonB-dependent receptor, a pectin methylesterase and a pectate lyase, as well as the TonB-dependent receptor, which is named *iroN* (ASF73_19920) and has been identified in the transposon screen (Figure S8).

We examined the ability of the gene deletion strains to degrade plant tissue and their impact on plant fresh weight during *rbohD* infection. With the exception of *gtf*, all other mutants showed phenotypes. Leaf degradation by the *dsbB* mutant was abolished in *rbohD*, similar as the *xps* mutant (Figure 5A). In accordance with the impaired leaf degradation, the *dsbB* mutant was also reduced in virulence as indicated by higher fresh weight of *dsbB* colonized *rbohD* plants (Figure 5B) and had a lower colonization level compared to wild type, similar to *xps* and *xpsxcs* mutants (Figure 5D). DsbB is involved in posttranslational modification of secreted enzymes, including proteins of the T2SS, which therefore explains the similar phenotypes between the mutants^38^. The *glucanase* and *iroN-glucanase* mutants showed reduced or delayed leaf degradation in *rbohD* (Figure 5A) and cell-free supernatant of liquid culture from the respective mutants revealed reduced degradation activities in *rbohD* leaf discs (Figure 5C). This finding suggests that the glucanase might be directly involved in polymer degradation. All mutants with reduced degradation activity were also reduced in overall virulence as indicated by higher fresh weight of *rbohD* plants (Figure 5B), while *glucanase* and *iroN-glucanase* mutants maintained wild type colonization levels (Figure 5D). The *glucanase* gene, which is absent in the *glucanase* and *iroN-glucanase* mutants (Figure S8), encodes a protein belonging to the glucanase superfamily (pfam13385) and contains a signal peptide for secretion. Remarkably, this *glucanase* mutant has a significant contribution to leaf degradation and virulence *in planta*.

**Figure 5.**
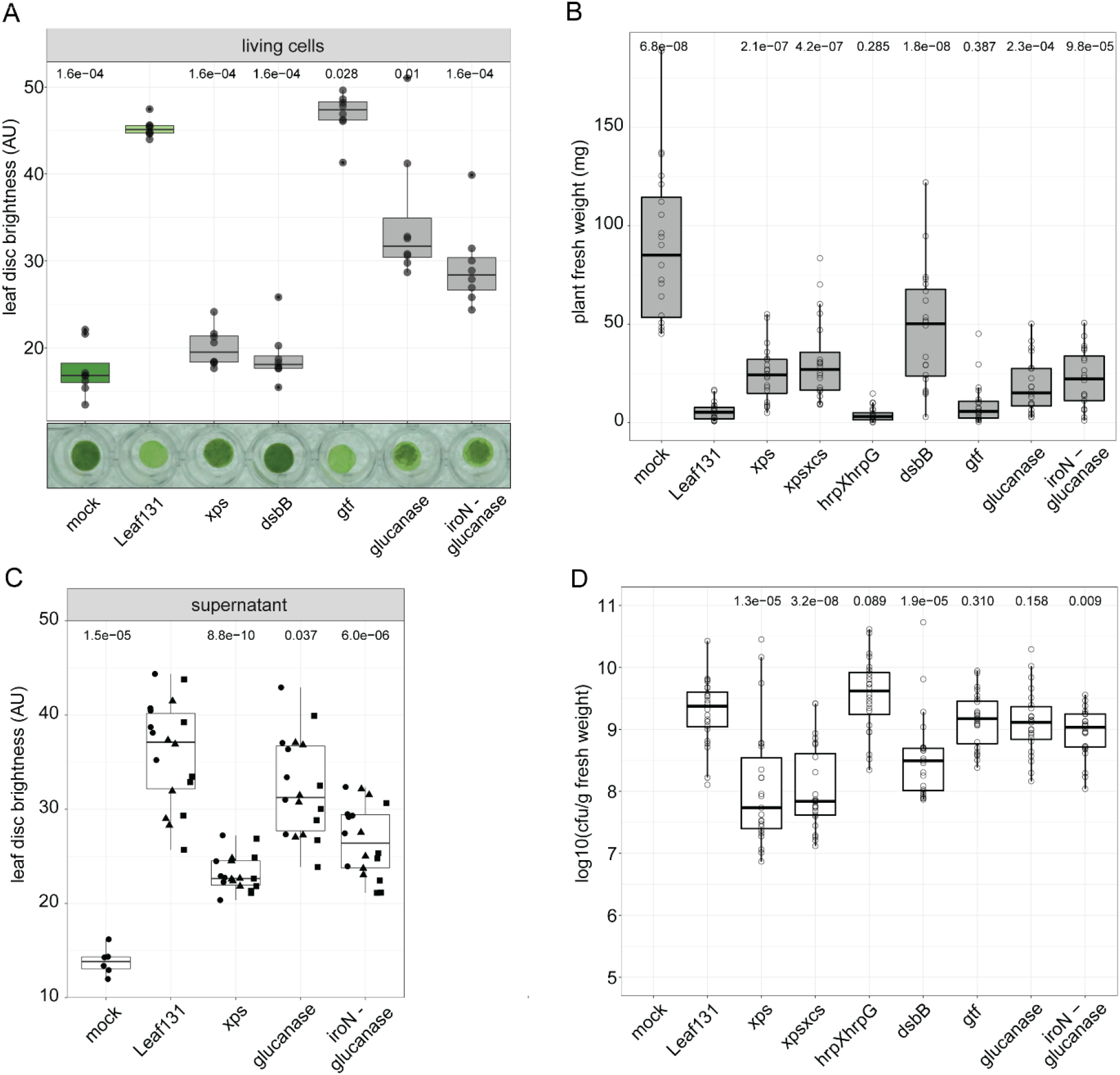
Additional virulence factors contribute to leaf degradation and virulence of *Xanthomonas* Leaf131. **A)** Leaf discs of five-week-old *rbohD* plants were mock (10 mM MgCl_2_) inoculated or with *Xanthomonas* Leaf131 wildtype (WT) or gene deletion mutants (OD=0.02) and incubated for 24 hours. **B)** Fresh weight of aboveground plant tissue of five-weeks-old gnotobiotic *rbohD* plants either mock inoculated or with *Xanthomonas* Leaf131 wildtype or gene deletion mutants. **C)** Leaf discs of five-week-old *rbohD* plants were mock treated (0.5x LB) or treated with cell-free supernatant (0.22 μm filter-sterilized) of liquid cultures from *Xanthomonas* Leaf131 wildtype and gene deletion mutants. Leaf discs were incubated for 24 hours at 22°C. Black circles, rectangles and squares indicate data from three bacterial cultures. **D)** Colony forming unit (CFU) counts of *Xanthomonas* Leaf131 per gram plant fresh weight from samples in B). Box plots show the median with upper and lower quartiles and whiskers present 1.5× interquartile range. Significant difference in A) (n=8), B) (n=20), C) (n=24) and D) (n=12) of gene deletion mutants compared to Leaf131 wildtype was determined by two-tailed Mann–Whitney *U*-test and p-values indicated above box plots.

### Virulence of *Xanthomonas* promotes abundance of specific community members

The secretion of extracellular enzymes is a crucial virulence factor of opportunistic *Xanthomonas* bacteria for plant colonization (Figure 3 and Figure 4), and our SynCom experiments revealed that plant disease and the microbiota shift in *rbohD* depends on the presence of *Xanthomonas* Leaf131 (Figure 1). To investigate whether both phenotypes are causally linked and dependent on T2SS-related virulence, we inoculated plants with the SynCom-137 and added either *Xanthomonas* Leaf131 wildtype or attenuated mutant strains. We determined the microbiota profiles by 16S rRNA amplicon sequencing and compared the community composition of the SynCom-137 containing *Xanthomonas* Leaf131 wildtype or mutants with the SynCom-137 without Leaf131 as a control. Quantification of the impact of each *Xanthomonas* strain on the community composition revealed a significant effect only of *Xanthomonas* Leaf131 wildtype (effect size 10.3%, p=0.0002), as observed previously (Figure 1A) but not the attenuated mutants in *rbohD* plants (Figure 6A). Consistently, only the presence of virulent *Xanthomonas* Leaf131, but not attenuated mutants with defective T2SS or *dsbB* knockout, increased the relative abundance of other commensals (Figure 6B). The addition of *Xanthomonas* Leaf131 wildtype to the SynCom-137 showed the characteristic shift in specific strains, namely *Sphingobium* Leaf26, *Xanthomonas* Leaf131, *Pseudomonas* strains Leaf58, Leaf127 and Leaf434 and *Sanguibacter* Leaf3 (Figure 6B and Figure S9A), as observed previously (Figure 1B). In contrast, inoculation of *rbohD* plants with SynCom-137 containing the attenuated *Xanthomonas* Leaf131 *xps, xpsxcs* or *dsbB* resulted in a similar overall community composition as the SynCom-137 alone in *rbohD* and in Col-0 plants, as indicated by few changes of individual strains in their relative abundance (Figure 6B and Figure S9B) and by overlapping clusters of the different conditions in a PCA (Figure S9C). In addition, the T2SS mutants showed reduced relative abundance, and *dsbB* was hardly detected by 16S rRNA amplicon sequencing (Figure 6C), which underlines the importance of these features for the competitiveness of *Xanthomonas* in the context of a bacterial community, similar to the plant inoculations with only *Xanthomonas* Leaf131 (Figure 4C).

**Figure 6.**
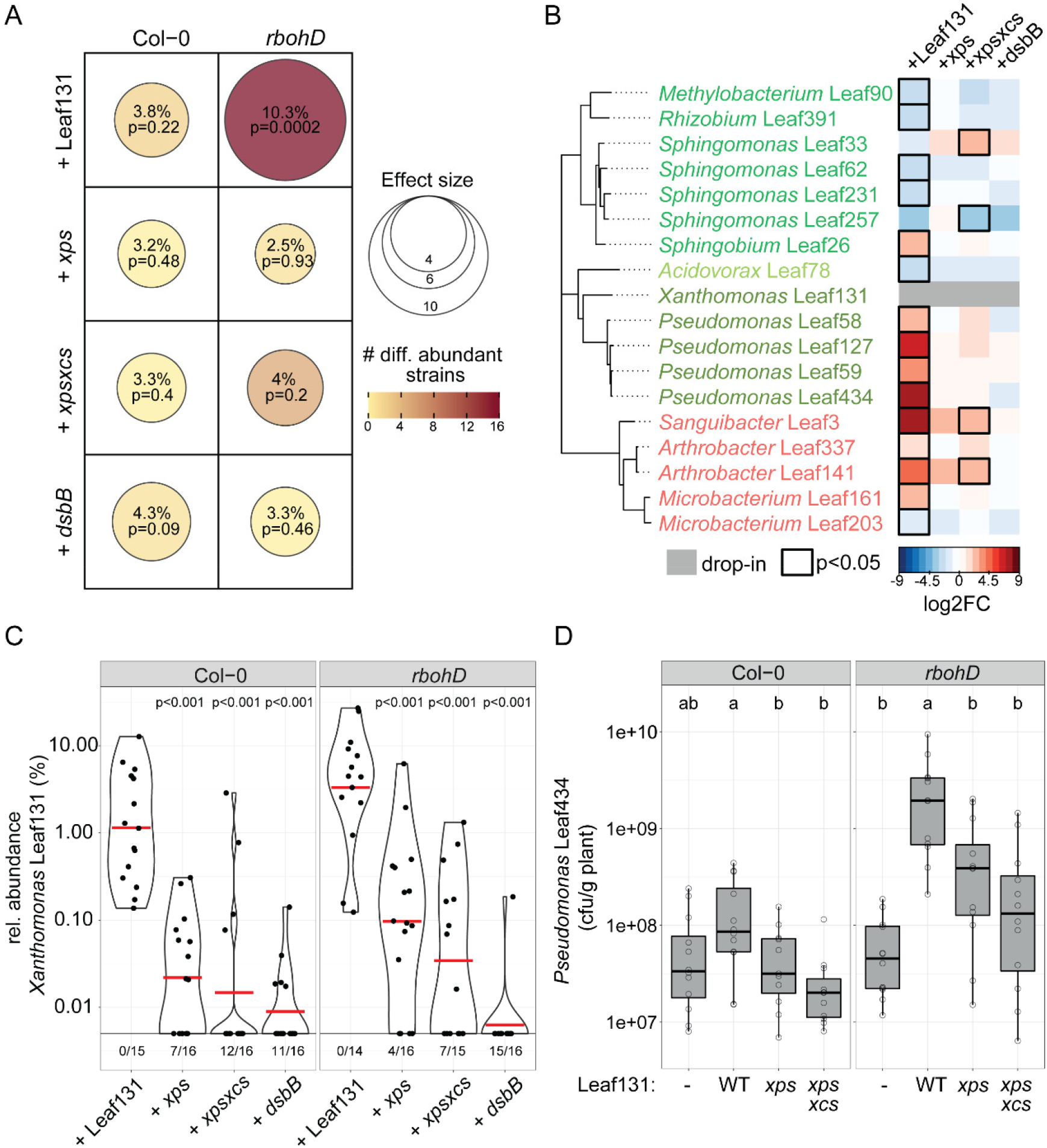
Microbiota shift in *rbohD* is dependent on T2SS-related virulence of *Xanthomonas* Leaf131. **A)** Composition of synthetic bacterial community SynCom-137 containing *Xanthomonas* Leaf131 wildtype or mutants *xps, xpsxcs*, and *dsbB* was compared to SynCom-137 alone in Col-0 and *rbohD* plants. Effect size represents percentage of total variance explained by genotype (shown by dot size and absolute value) and statistical significance is expressed with p-values determined by PERMANOVA (Benjamini–Hochberg adjusted, n = 16). Number of differentially abundant strains (as shown in panel A) is represented by dot colour. B) Heatmap shows subset of strains of SynCom-137 with significant log_2_ fold changes (log2FC, p-value < 0.05) in *rbohD* plants inoculated either with only SynCom-137 or with SynCom-137 containing *Xanthomonas* Leaf131 wildtype or the mutants *xps, xpsxcs*, and *dsbB*. Black rectangles show significant changes, p-value < 0.05 (n = 16, Wald test, Benjamini–Hochberg adjusted). Complete heatmap of all strains in SynCom-137 is shown in Figure S8B. **C)** Relative abundance of *Xanthomonas* Leaf131 wildtype or the mutants *xps, xpsxcs*, and *dsbB* within SynCom-137 in Col-0 and *rbohD* plants. Ratio below violin plots represent frequency of samples where *Xanthomonas* Leaf131 was not detected. Violin plots show the median with upper and lower quartiles (n = 16, two-tailed Mann–Whitney *U*-test, p-values indicated above box plots). **D)** Colony forming unit (CFU) counts of *Pseudomonas* Leaf434 per gram plant fresh weight after inoculation of germ-free Col-0 and *rbohD* plants with *Pseudomonas* Leaf434 as single inoculation (-) or as binary inoculation with either *Xanthomonas* Leaf131 wildtype or the mutants *xps* and *xpsxcs*. Box plots show the median with upper and lower quartiles and whiskers present 1.5× interquartile range (n = 12, two-tailed Mann–Whitney *U*-test, p-values indicated above box plots).

Furthermore, we examined in a binary strain inoculation experiment the colonization level of the commensal *Pseudomonas* Leaf434 in response to attenuated *Xanthomonas* Leaf131. Strikingly, the increase in the commensal *Pseudomonas* Leaf434 observed during co-inoculation with virulent *Xanthomonas* Leaf131 was significantly reduced when the T2SS mutants *xps* and *xpsxcs* were paired with the *Pseudomonas* strain (Figure 6D). This finding supports the conclusion that commensals are enriched in their abundance in plants due to the virulence of *Xanthomonas* Leaf131, which is particularly pronounced in immunocompromized *rbohD* plants.

In summary, our results suggest that specific microbiota members benefit indirectly from *Xanthomonas* Leaf131 due to T2SS-dependent virulence causing plant disease, rather than from the presence of *Xanthomonas* Leaf131 or the knockout of RBOHD *per se*.

## Discussion

Host-associated microbial communities play a central role for the health of animals and plants^3,39^. In humans, dysbiosis is associated with many pathologies such as inflammatory bowel disease, diabetes, and allergies, and is often accompanied by a decrease in microbial diversity, the presence of potentially harmful microbes, or absence of beneficial microbes^4,5^. However, the concept of dysbiosis is controversial because the causal relationships are often unclear, e.g. whether the observed changes in the microbiota are caused by the host genotype or by infection with a pathogen, and whether a shift in the microbial composition is the consequence of host disease or promotes disease^4,40^. The composition of the microbiota also depends on the diet, among other environmental factors, making it challenging to define a balanced microbiota in a healthy state^3,41^. Therefore, dysbiosis is considered a condition with distorted microbiota with various compositional states – but always associated to disease and often characterized by weakening of host control over microbial growth^42,43^. Studies involving germ-free animal models with defined microbiota are increasingly being used to help discern cause and consequences^44,45^ with some inherent limitations^46^.

For plants, several recent experimental studies described that a functional immune system is required to maintain microbiota homeostasis and prevent dysbiosis^9,10,47^. Our previous finding that *A. thaliana rbohD* mutants display a microbiota shift and the identification of *Xanthomonas* Leaf131 and Leaf148 as opportunistic pathogens^9^ gave us the opportunity to disentangle the causation of dysbiosis in the representative *At*-LSPHERE leaf microbiota. We found that T2SS-dependent virulence of *Xanthomonas* leads to a) tissue damage in *rbohD*, b) increased relative abundance of *Xanthomonas* Leaf131 and c) was responsible for the shift in the composition of the residual leaf microbiota resulting in dysbiosis. Because *Xanthomonas* Leaf131 was an inherent constituent of the microbiota, the conditional pathogenicity was unlocked by *rbohD*, which results in a dysbiotic microbial community characterized by increased abundance of *Xanthomonas* Leaf131 and other strains (Figure 1 and Figure 6). While several studies have reported dysbiosis in the phyllosphere of plants infected with a pathogen^48-52^, it remains to be shown whether the pathogen invaded the microbial community as external agent or was part of the microbiota and was kept under control. Indeed, environmental conditions and protective microbiota members determine the virulence of pathogens^25,53-56^.

Our results of the SynCom experiments in Col-0 and *rbohD* with virulent and virulence attenuated *Xanthomonas* suggest that the enrichment of selected commensal strains (e.g. *Pseudomonas* Leaf434) within the bacterial community is due to *Xanthomonas* Leaf131-induced disease. Due to the leaf tissue degradation caused by *Xanthomonas* Leaf131, these commensals might benefit from nutrients released from the plant as “public good” and their metabolic capacity and/or faster growth rate allow them to reach a higher population size within the overall microbiota. Metabolic interactions in the phyllosphere shape the microbiota by making simple sugars available or complementing vitamin auxotrophies^57,58^. To track the bacterial behaviour by transcriptomic or proteomic community profiling could provide further insights into the microbial interactions within the microbiota.

We found that leaf tissue damage caused by secreted CWDE via the T2SS Xps is a major virulence strategy of *Xanthomonas* Leaf131 and Leaf148 during infection of *A. thaliana*. Consequently, *xps* deletion mutants did not impact growth of Col-0 wildtype and have a reduced impact on *rbohD* plants. However, it is still unclear what underlies the context-dependent pathogenicity of *Xanthomonas* Leaf131 and Leaf148 in *rbohD* plants. Virulence in these opportunistic *Xanthomonas* strains could be regulated by *rbohD*-specific cues (e.g. nutrients, signalling molecules or absence of ROS) that trigger a behavioural switch in *Xanthomonas* towards a pathogenic lifestyle. It was recently shown in *X. citri* pv. *citri* that degradation products from the plant cell wall polymer xyloglucan induces transcription of virulence factors^59^. In our leaf degradation experiments using supernatants, *Xanthomonas* produced CWDE during incubation in rich media, suggesting that CWDE production is a constitutive trait and might not be dependent on host signals, although at this time we cannot exclude that expression levels are enhanced *in planta*. In this context it is, however, interesting to note that the HrpG and HrpX master regulators of virulence – as described in multiple *Xanthomonas* pathogens mainly for T3SS-mediated virulence^32,34,60,61^ – were not required for tissue degradation activity and virulence of *Xanthomonas* Leaf131 (Figure S4). Transcriptomic studies in *Xanthomonas campestris* pv. *campestris* found that the T2SS genes and some (but not all) T2SS substrates were regulated by HrpG and that only a small subset of all genes regulated were part of the HrpG-regulon, at least during hydathode infection of cauliflower^31,60^, which could explain the absence of a phenotype for *hrpGhrpX* knockout in *Xanthomonas* Leaf131. We identified several T2SS-dependent proteins to be secreted by *Xanthomonas* Leaf131 and Leaf148 in liquid culture. Interestingly, homologous proteins of an endoglucanase (ASF73_13775) have also been detected in supernatant of *X. campestris* pv. *vesicatoria*^36^ and of *X. oryzae* pv. *orzyae*^29^, where the protein was required for virulence in rice for the latter. In addition, we identified a serine protease (ASF73_18370) that was detected in supernatants of Leaf131 rather than of the T2SS mutant. A homologue of this enzyme was described to be required for growth *in planta* of *X. citri* pv. *citri*^62^ and to be secreted by *X. campestris* pv. *vesicatoria*, but in a T2SS-independent manner^31^. We have not observed a reduction in leaf tissue degradation in the gene knockout strains for the endoglucanase or serine protease, which is not surprising given that a deletion of a single T2SS substrate often does not show a phenotype presumably due to functional redundancy among the secreted proteins^29,31^. Overall, our results match the observations from other studies that the importance of the HrpG and HrpX regulators and the T2SS substrates contributing to virulence can differ between *Xanthomonas* bacteria^31,63^.

Context-dependent pathogenicity of opportunistic *Xanthomonas* strains might rely on plant susceptibility due to altered immune signalling or physical barriers. The plant immune system detects microbial activity and monitors the cell wall integrity^64,65^. Loss of MAMP/DAMP-induced ROS production by RBOHD results in impaired immune signalling and increased susceptibility to bacterial and fungal pathogens^14,66,67^ as well as reduced cell wall remodelling and lignification^16,17,68^. To explain susceptibility to opportunistic *Xanthomonas, rbohD* plants could mount an insufficient defence response. Many pathogens secrete CWDE to degrade plant polymers at certain stages during the infection process^65,69^ and, in turn, defects in cell wall composition make plants more susceptible^70^. As such, *rbohD* plants might have cell wall defects due to altered polymer crosslinking, which is in accordance with our data showing that tissue degradation activity of cell-free supernatant is higher in *rbohD* compared to Col-0 leaf discs. In that case, opportunistic *Xanthomonas* would secrete CWDE that break down a vulnerable (pre-formed) cell wall of *rbohD* plants. Strikingly, we have identified a single gene, *glucanase* (ASF73_19940) that is required for full leaf tissue degradation and virulence in *rbohD* plants, and encodes a protein annotated with a secretion signal and a glucanase/concanavalin A-like lectin domain, which is potentially involved in carbohydrate processing or adhesion. Mammalian NADPH oxidases produce ROS as cell-to-cell messenger regulating the intestinal barrier which is required for microbiota homeostasis^71^ and ROS also form a physical barrier, which is thought to keep certain bacteria at distance from the epithelial surface^21,72^. This draws attention to striking similarities in the molecular mechanisms for host control of microbiota homeostasis across animal and plant kingdoms.

In conclusion, our study revealed the importance of the T2SS for opportunistic *Xanthomonas* strains both for their interaction with the plant and for their competitiveness within the microbiota. The conditional pathogenicity of this opportunistic microbiota member depends on the host genotype and impacts both plant health and the microbial community. Our findings establish a causal link between a single plant gene to a specific genus of bacteria that drives a microbiota shift and highlight the crucial role of opportunistic pathogens in dysbiosis.

## Materials and Methods

### Plant growth conditions in soil

*A. thaliana* wildtype Col-0, *bak1/bkk1/cerk1* (*bbc*)^53^, *fls2/efr/cerk1* (*fec*)^73^, *rbohD* knockout mutant^14^ and complementation line *rbohD/pRBOHD::RBOHD-FLAG* (*rbohD/RBOHD*)^66^ were used in this study.

*A. thaliana* plants for leaf degradation assays were grown in peat-based potting soil (Substrat 1, Klasmann-Deilmann, Germany) in a growth chamber (CU-41L4, Percival) under controlled conditions (11-h light cycle, 22° C, 65% relative humidity, light intensity (PAR) 200 μmol*s^-1^*cm^-2^). Seeds were treated with 70% ethanol for two minutes, sown on soil and stratified for two days at 4° C in the dark.

### Gnotobiotic plant growth and bacterial inoculation

Gnotobiotic plants were prepared and grown as described previously^9^. Briefly, surface sterilized seeds were placed in sterile microboxes filled with calcined clay (Diamond Pro Calcined Clay Drying Agent) and supplemented with 0.5x Murashige and Skoog (MS) medium including vitamins, pH 5.8 (M0222.0050, Duchefa). Plants were grown in growth chambers (CU-41L4, Percival) set to 22 °C and 54% relative humidity and run under an 11-h light cycle as reported previously^9^.

For the SynCom, 138 strains were selected on the basis of the *At*-LSPHERE strain collection (Supplementary table 5) to have maximal phylogenetic diversity and to distinguish all strains with 100% sequence identity representing amplicon sequence variants (ASVs) by amplifying the V5-V7 region of the 16S rDNA gene amplified with the primers 799F and 1193R, as described previously^9^. In addition, *Xanthomonas* sp. Leaf131 was used as single inocula or mixed into the SynCom-137. *Xanthomonas* sp. Leaf148 has identical V5-V7 region of the 16S rDNA gene compared to *Xanthomonas* sp. Leaf131 and 87% average nucleotide identity (ANI) across the entire draft genome^9^.

Bacterial growth, mixing of the synthetic community and plant inoculation were done as described before^9^. Briefly, bacteria were grown on R2A-agar (Sigma-Aldrich) plates containing 0.5% v/v methanol (R2A-MeOH) for 4 d at 22 °C, restreaked on fresh R2A-MeOH plates and grown for 3 d. To prepare the SynCom mix, similar volume of cells for each strain were scraped from agar plates and resuspended individually in 10 mM MgCl_2_. Each strain was mixed in equal volume ratio for inoculum mix. Germ-free, 11-days-old seedlings were inoculated with 200 μL bacterial solution. Plants were harvested between 35 to 38 days after germination. Experiments with SynCom, single strain or binary strain inocula were done in same procedure.

To determine bacterial colonization levels, the plant phyllosphere was harvested, weighed and homogenized in 10 mM MgCl_2_ and a dilution series plated on R2A-MeOH agar plates to count cfu after 2 days incubation at 28 °C. To extract DNA for 16S rRNA amplicon sequencing, the plant phyllosphere was harvested, weighed and stored at -80 °C.

### DNA extraction and 16S rRNA amplicon sequencing

DNA extraction and 16S rRNA amplicon sequencing was done according to the procedure that was previously published^9^. Briefly, plants were frozen in sampling tubes for DNA extraction using the FastDNA SPIN Kit for Soil (MP Biomedicals). Samples were freeze-dried (Alpha 2-4 LD Plus, Christ) and ground to powder at 30 Hz for 2 × 45 s (TissueLyzer II, Qiagen). DNA was extracted according to the manufacturers instructions. DNA was quantified using dsDNA QuantiFluor (Promega) and normalized to 1 ng*μL^−1^ in low-binding 96-well plates. The 16S rDNA amplicon library was generated as previously described^9^, but PCR reactions were not done in triplicate here. A first PCR reaction using 799F and 1193R using Taq polymerase (Bioron DSF Taq) followed by a clean up using Antarctic phosphatase (New England Biolabs, NEB) and exonuclease I (NEB) to digest unused primers and a second barcoding PCR reactions.

The 16S rRNA band intensity of each sample after barcoding PCR was visually inspected on an 2% agarose gel and samples from one plate were pooled together at approximately equal ratio. The pooled DNA was cleaned up with AMPure magnetic beads using 1:1 boead-to-DNA ratio. The cleaned-up DNA was run on a 2.5% agarose gel to separate bacterial 16S rRNA amplicon band from plastid 16S rRNA amplicon and excised from the gel. The DNA was extracted from the gel and all pools were combined in equal ratio of each sample in the final pool. The pooled 16S rRNA amplicon library was cleaned twice with AMPure magnetic beads using a bead-to-DNA ratio of 0.7 to remove small DNA fragments according to the manufacturer’s protocol. The amplicon length distribution of the library was assessed on a 2200 TapeStation using HS D1000 (Agilent), resulting in an effective library size of 560–620 bp. Sequencing was performed on a MiSeq desktop sequencer (Illumina) at the Genetic Diversity Centre Zurich using the MiSeq reagent kit v.3 (paired end, 2 × 300 bp). Denaturation, dilution and addition of 12% PhiX to the DNA library were performed according to the manufacturer’s instructions. Custom sequencing primers were used as described previously^9^.

### Data analysis of 16S amplicon sequencing

16S rRNA amplicon data processing was done as described previously^9,23^. Briefly, a database was created of each strain in the SynCom represented by one ASV reference. Paired-end sequencing reads were merged using the USEARCH v.11.0.667-i86 linux64 command fastq_mergepairs^74^, with a minimum overlap of 16 bp and a minimum identity of 90%. Merged reads were quality filtered using - fastq_filter with a maximum expected error of one and a minimum length of 200 bp. The reads were classified and counted using -otutab with 100% identity to 16S rRNA gene reference sequences, which resulted in an ASV count table (Supplementary table 5).

The ASV table of each experiment was processed in R v.3.6.3 as described previously^9^. To account for varying sequencing depths between samples, the ASV table was log-normalized and variance-stabilized by DESeq2 v.1.14.1. To examine the effect on individual strains between the test and control conditions, the output of DESeq2 provided log2 fold change values and strains were considered to be differentially abundant according to Wald test implemented in DESeq2. p-values were adjusted for multiple testing using the Benjamini-Hochberg (BH) method implemented in DESeq2. The differential strain abundances between the test and control conditions were visualized as heatmap. To assess the overall effect on communities, PCA was performed with the transformed ASV table using the prcomp command. The effect size represents the variance explained by the compared factor and was calculated on Euclidean distances followed by a PERMANOVA to test for statistical significance using the adonis command of the package vegan v.2 v.5-4. To summarize the relative abundance of *Xanthomonas* Leaf131 in a sample, the relative abundance values were calculated by proportional normalization of each sample by its sequencing depth.

The following R packages were used during analysis and visualization: ape v.5.4 (ref.^75^), ggplot2 v.3.3.0 (ref.^76^, vegan v.2.5-4 (ref.^77^) and DESeq2 v.1.14.1 (ref.^78^). ggpubr v.0.3.0 (ref.^79^).

### Transposon mutagenesis screen

The transposon (Tn) mutagenesis screen was done by transforming electrocompetent cells of Xanthomonas Leaf131 with the pAK415 plasmid containing a vanillate-inducible, hyperactive transposase Himar1C9W^37^. After electroporation, bacteria were recovered in 500 μL LB containing 500 μM vanillate at 28°C shaking for 4 hours to induce temporal expression of the transposase.

Single colonies were selected on LB-agar plates containing 50 μg*mL^-1^ kanamycin grown for two days at 28°C. Colonies were picked and grown in 100 μL liquid LB medium in 96-well plates. After measuring OD_600_ to monitor bacterial growth, each transposon mutant colony was tested for its potential to degrade a leaf disc from five-week-old *A. thaliana rbohD* plants. In total, 6016 mutants were screened and 92 Tn mutants showed either delayed or no leaf degradation after 24 hours (Supplementary table 2). To validate the phenotype, we retested all identified Tn mutants on eight leaf discs of *rbohD* plants. For this, bacteria were grown on R2A-MeOH agar plates and resuspended in 10 mM MgCl_2_ and leaf discs inoculated at an OD_600_ of 0.02. After the validation screening, 34 Tn mutants were confirmed to have a deficiency in *rbohD* leaf degradation after 48 hours.

To identify the Tn insertion site for each candidate Tn mutant, we mapped the genomic location by a nested semi-degenerated primer PCR approach^37,80^. The first PCR reaction was done with a mixture of semi-degenerated primers (Arb-P1, Arb-P2, Arb-P3) mixed in equal ratio and a transposon-specific primer (pAK411_nested1). In a 25 μL PCR reaction, 1 μL of 10 μM forward and reverse primers, 0.2 μL Phusion polymerase, 5 μL of 5x Phusion GC buffer, 1.25 μL dNTP mix, 0.75 μL DMSO and 1 μL of supernatant from heat-killed bacterial suspension were used as template. Hot-start PCR was done in a thermocycler using the following settings: initial 98°C for 3 min, 10 cycles of 98°C for 10 seconds, an annealing step starting at 35°C for 20 seconds and increasing 1°C in each cycle until 45°C, elongation at 72°C for 30 seconds; the temperature gradient cycles were followed by 25 cycles of 98°C for 10 seconds, annealing at 45°C for 20 seconds, elongation at 72°C for 30 seconds and a final elongation at 72°C for 5 minutes. The PCR product was visually inspected on an agarose gel and cleaned up using NucleoSpin gel and PCR clean-up kit (Machery-Nagel, Düren, Germany). The DNA fragments from the first PCR were used as templates in a second PCR using sequence-specific (nested) primer binding on the transposon and on a sequence introduced by the semi-degenerated primers. The PCR reaction was done in 50 μL total volume using 2 μL of 10 μM Anchor-P primer and pAK411_nested2 primer, 0.4 μL Phusion polymerase, 10 μL of 5x Phusion GC buffer, 2.5 μL dNTP mix, 1.5 μL DMSO and 1 μL of purified PCR product as template. Hot-start PCR was done with following settings: initial denaturing at 98°C for 3 min, 5 cycles of 98°C for 10 seconds, an annealing step starting at 63°C for 20 seconds and decreasing 1°C in each cycle until 59°C, elongation at 72°C for 30 seconds; the temperature gradient cycles were followed by 30 cycles of 98°C for 10 seconds, annealing at 58°C for 20 seconds, elongation at 72°C for 30 seconds and a final elongation at 72°C for 5 minutes. The product of the second PCR was cleaned-up and Sanger sequenced using primer pAK411_nested2. The genomic insertion site was identified by using the PCR amplified sequence in a BLAST search against the genome of *Xanthomonas* sp. Leaf131 (NCBI:txid1736270) at NCBI and IMG/JGI.

### Leaf disc degradation assay

Leaf discs of 5- or 6-weeks-old *A. thaliana* plants grown in soil were collected using a 4 mm diameter biopsy puncher (BPP-40F, kai medical) and placed with the adaxial side up in a clear flat-bottom 96-well plate (655101, Greiner bio-one) filled with 90 μL milli-Q purified (mQ) water. *Xanthomonas* were grown on R2A-MeOH agar plates for 2 days at 22°C; bacterial cells were scraped off, resuspended in 10 mM MgCl_2_ by vortexing for 2 min and the bacterial solution was adjusted to OD_600_ = 0.1. Leaf discs were inoculated with 10 μL of bacterial suspension and incubated at 22°C for up to 48 hours in the dark. Digital images were taken at regular intervals under standardized conditions using a black box and a light screen illuminating leaf discs from below to monitor leaf tissue degradation.

### Quantification of leaf disc brightness

To quantify leaf tissue degradation, we developed a computational script MatlabR2022a (MathWorks, USA), which recognizes leaf discs in a 96-well plate, measures surface area, brightness of the red channel (in RGB images) and computes a ‘roughness’ parameter.

In short: The script normalizes the brightness of the images using the ‘illumgray’ and ‘chromadapt’ functions implemented in Matlab. Subsequently a binary mask is created, separating the area occupied by leaf discs from the rest of the image. Discs that deviate in ‘roundness’ are discarded from the analysis since they are likely to be broken or folded. The roughness parameter is created using the ‘Sobel’ edge detection function on the isolated discs in the red channel and computing the total number of pixels recognized as edge within each disc. The area of each disc is computed by counting the number of pixels per disc times the pixel size retrieved from an image scaling step. The brightness value represents the mean brightness value of the red channel for each individual leaf disc.

The MatlabR2022a code and user manual is available at: https://github.com/gaebeleinC/leaf-disc_quantification

### Transformation of electrocompetent *Xanthomonas* cells

Electrocompetent *Xanthomonas* cells were made by an established protocol^81^. Exponentially growing *Xanthomonas* cells in 200 mL LB at 28°C with an OD_600_ between 0.6 and 1 were cooled on ice for 20 min and kept on ice for the entire procedure. Cells were collected by centrifugation for 15 min at 4000xg at 4°C and washed three times in chilled, sterile 10% glycerol to remove growth medium. After the final washing step, cells were concentrated approximately 100-fold compared to initial volume in 10% glycerol and aliquots frozen at -80°C.

Electrocompetent *Xanthomonas* cells were thawn on ice and 50 μL mixed with 200 ng plasmid. Cells were transformed by electroporation in 1 mm electro-cuvettes applying 1.8 kV electric current. Cells were recovered in LB medium for 2-4 hours shaking at 28°C before plating 100 μL on LB-agar plates containing selective antibiotics.

### Bacterial gene knockout strains

Markerless gene deletion in *Xanthomonas* strains (Supplementary table 3) were made according to a method based on double homologous recombination using the suicide plasmid pK18mobSacB as vector^82^. Gene deletion plasmids were designed to result in in-frame deletion of the gene of interest while leaving an open reading frame of 3-4 amino acid peptide. Briefly, 500 bp of flanking regions upstream and downstream of the gene of interest were amplified by PCR and cloned into pK18mobSacB plasmid. The plasmids were cloned using either classical restriction enzyme digest or Gibson Assembly (primer list in Supplementary table 4) in *E. coli* Dh5α. Gene deletion constructs were confirmed by Sanger sequencing.

Electrocompetent *Xanthomonas* cells were transformed, recovered in LB medium for 2-4 hours and transformed cells were selected on LB-agar plates containing 50 μg*mL^-1^ kanamycin. Transformed cells were re-streaked on fresh selective LB-agar plates and a single colony resuspended in LB medium for 2 hours before plating on LB agar plates containing 5% sucrose to select for double cross-over events due to homologous recombination and chromosomal deletion of the gene of interest and the vector backbone. After sucrose selection, individual colonies were tested for sensitivity to kanamycin. Cells were re-streaked to obtain single colonies that were cultured and frozen in 25% glycerol at -80°C. Genomic deletion was confirmed by PCR using primers outside of flanking regions and Sanger sequencing the PCR product and by the absence of PCR product using primers inside genomic deletion.

### Bacterial supernatant of liquid culture

*Xanthomonas* were grown in triplicates in 100 mL liquid 0.5xLB medium until late exponential growth phase (approx. OD_600_ = 2) at 28°C shaking. Cells were harvested by centrifugation at 4000x g for 15 min and washed twice in 10 mM MgCl_2_. Bacterial cells were resuspended in 10-20 mL fresh 0.5x LB medium at an OD_600_ = 3 and incubated for 4h in flasks at 28°C shaking. To obtain the cell-free supernatant, we centrifuged the samples at 4000x g for 15 min to remove bacteria and filter-sterilized the supernatant using 0.22 μm filter units (#99505, „rapid”-Filtermax, TTP, Trasadingen, Switzerland) and a vacuum pump. 10 mL of Cell-free supernatant was concentrated 10-fold by using Ultrafiltration Units Amicon-15 with a molecular weight cut-off 10 kDa (Merck, Darmstadt, Germany) and centrifugation at 3500 xg at 4°C for 20 to 40 min. Cell-free supernatants were directly tested for leaf degradation activity and kept on ice until further processing for protein analysis.

Cell-free supernatant or concentrated supernatant was applied to leaf discs to test for tissue degradation activity. Leaf discs were collected from five- or six-week-old plants and floated in 40 μL mQ water in 96-well plate. To each leaf disc 40 μL supernatant was added. Leaf discs were incubated at 22°C and photographs taken at regular intervals.

### Analysis of protein bands by LC-MS/MS

To test for the secretion of proteins, the concentrated, cell-free supernatant of *Xanthomonas* liquid cultures was obtained as described above. Protein concentration of the concentrated supernatant was determined by the Pierce BCA assay kit (ThermoFischer Scientific, Reinach, Switzerland) according to the manufactures instructions. Protein content of supernatant samples were normalized and analyzed using SDS-PAGE (mPAGE Bis-Tris 8%, Merck, Darmstadt, Germany) revealing specific protein bands in the supernatant while comparing wildtype and the T2SS mutant (*xpsxcs*) of *Xanthomonas* Leaf131 and Leaf148. The protein bands of interest were cut-out and identified by in-gel digestion and LC-MS/MS analysis as described previously^83^.

### Substrate degradation by secreted enzymes in agar plates

Agar plate assays to detect glucanase, xylanase, pectate lyase and polygalacturonase or protease activity were modified after refs.^84-86^.

*Xanthomonas* strains were streaked on R2A-MeOH plates and grown at 22°C for two days. Bacterial cells were scraped off and resuspended in 1 mL 10 mM MgCl_2_ by vortexing for 5 min to disperse cell aggregates. Cell density was adjusted to OD_600_ = 0.4 and 4 μL of the bacterial suspensions were spotted on R2A agar plates either containing 0.5% sodium carboxymethyl cellulose (CMC; Sigma-Aldrich, C5678), 0.05% Remazol Brilliant Blue-Xylan (RBB-Xylan; Sigma-Aldrich, M5019), 0.1% Azo-Xyloglucan (Megazyme, S-AZXG) or 0.1% polygalacturonic acid in 1M sodium phosphate buffer pH 7.0 (PGA; Sigma-Aldrich, 81325) or on 1.5% agar plates containing 3% skimmed milk powder (Rapilait), 1% peptone, 0.025% MgSO_4_ and 0.05% K_2_HPO_4_, respectively. The plates were incubated at 22°C and photographs were taken at regular intervals.

Glucanase activity can be detected by yellow halos against the red background after staining with 0.1% Congo Red (Sigma-Aldrich, C6767) dye solution (solved in 50% ethanol) for 30 min and destaining with 1 M NaCl for 15 min. Pectate lyase or polygalacturonase activity can be detected by light pink halos against the darker pink background after staining with 0.05% Ruthenium Red (Sigma-Aldrich, R2751) dye solution (solved in water) for 30 min and destaining with water. Xylanase or protease activity can be detected by a light blue or clear halo forming around the colonies, respectively.

## Data and code availability

Customized code to analyse data and generate figures can be found at https://github.com/MicrobiologyETHZ/phylloR. Raw data of 16S rRNA amplicon sequencing will be made available at the European Nucleotide Archive upon peer-reviewed publication.

## Acknowledgements

The authors would like to thank Gabriella Petti, Clément Lefebvre and Trisha Steward for experimental help, Christopher Field for bioinformatic support and Martin Schäfer for fruitful discussions. Silvia Kobel from the Genetic Diversity Centre Zurich for assistance in library preparation and sequencing. The authors acknowledge funding from the German Research Foundation (DECRyPT, no. SPP2125), from the ETH Zurich Foundation (Career Seed Award), and the NCCR Microbiomes, funded by the Swiss National Science Foundation.

