## Supplementary Figures for "Dysbiosis of a leaf microbiome is caused by enzyme secretion of opportunistic *Xanthomonas* strains"

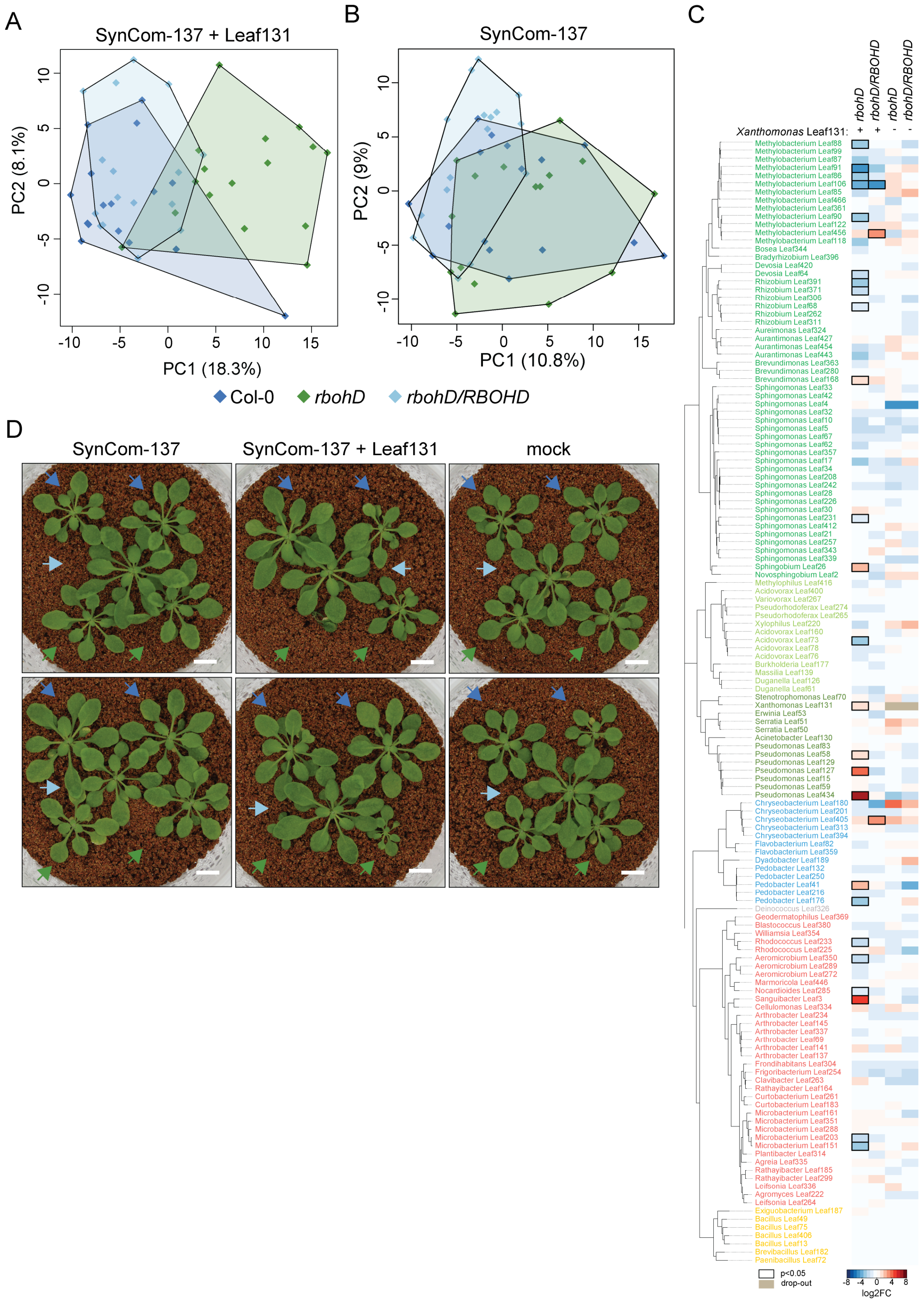


**Figure S1. Microbiota composition of SynCom-137 and plant phenotype in presence and absence of *Xanthomonas* Leaf131. A)** Principal component analysis of SynCom-137+*Xanthomonas* Leaf131 and of **B)** SynCom-137 in Col-0 (blue), *rbohD* (green) and *rbohD/RBOHD* (light blue). Axes show principal components PC1 and PC2 with their explained variance (%). **C)** Heatmap shows $log_2$ fold changes (log2FC) of strains in SynCom-137 in *rbohD* or *rbohD*/*RBOHD* compared to Col-0 wild-type plants in the presence (+) or absence (-) of *Xanthomonas* Leaf131. Black rectangles show significant changes, p-value < 0.05 (n = 16, Wald test, Benjamini–Hochberg adjusted). **D)** Plant phenotype of Col-0 (blue arrow), *rbohD* (green arrow) and *rbohD/RBOHD* (light blue arrow) mock inoculated or with SynCom-137 or SynCom-137+*Xanthomonas* Leaf131.

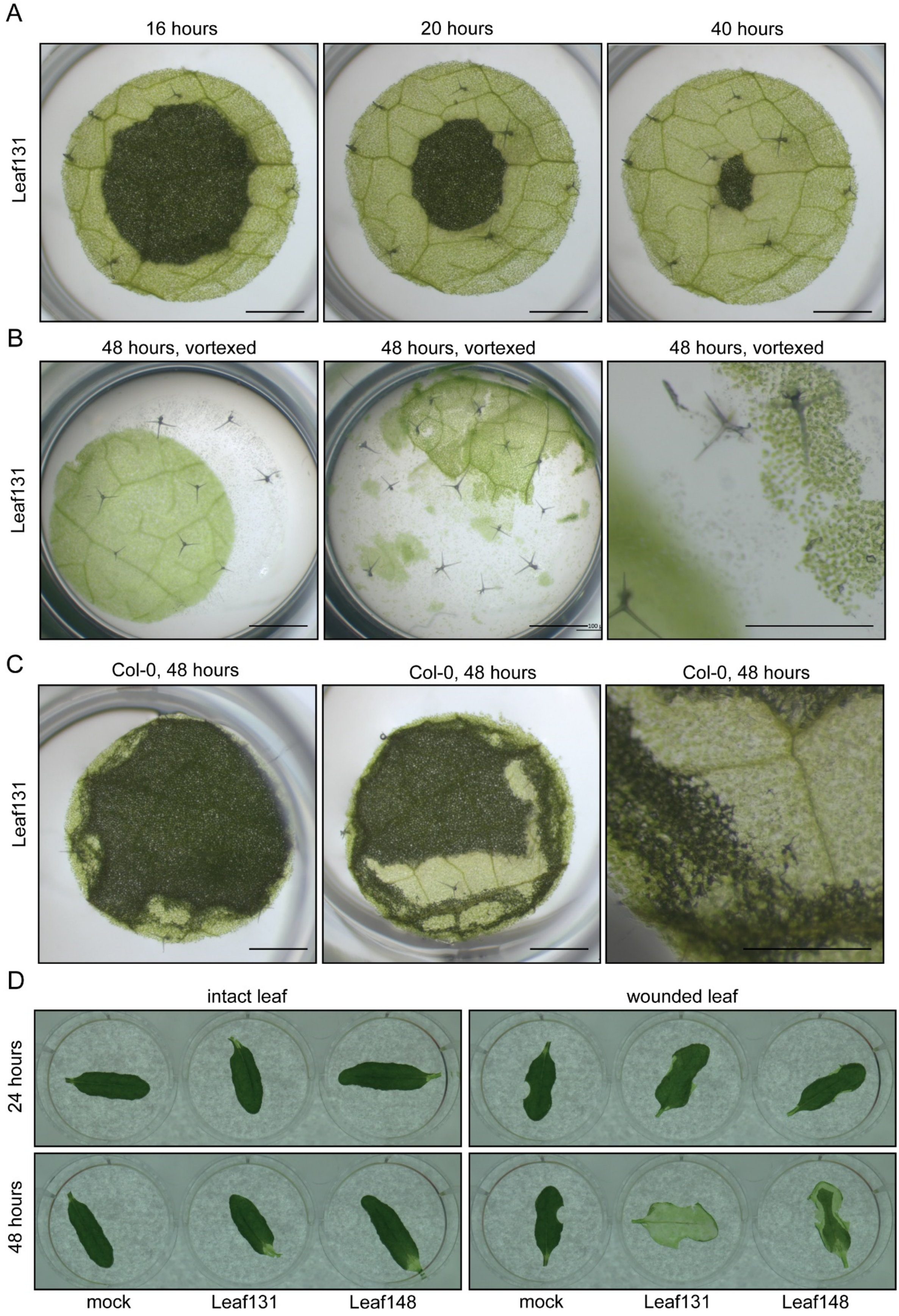


**Figure S2. *Xanthomonas* disrupts leaf tissue cohesion and requires plant wound. A)** Time-course of leaf discs from five-week-old *rbohD* plants inoculated with *Xanthomonas* Leaf131 (OD=0.02).

Scale bar represents 1 mm. **B)** Leaf discs of *rbohD* plants in 96-well plate after 48 hours incubation with *Xanthomonas* Leaf131 were vortexed for two seconds. Scale bar represents 1 mm in left and middle panel, and 0.5 mm in right panel. **C)** Leaf discs of five-week-old Col-0 plants after 48 hours incubation with *Xanthomonas* Leaf131. Scale bar represents 1 mm in left and middle panel, and 0.5 mm in right panel. **D)** Leaves of five-week-old rbohD plants floating in water with undamaged leaf edge (left panels) or wounded leaf edge (right panels) were mock inoculated (10 mM $MgCl_2$) or with *Xanthomonas* Leaf131 or Leaf148 (OD=0.02).

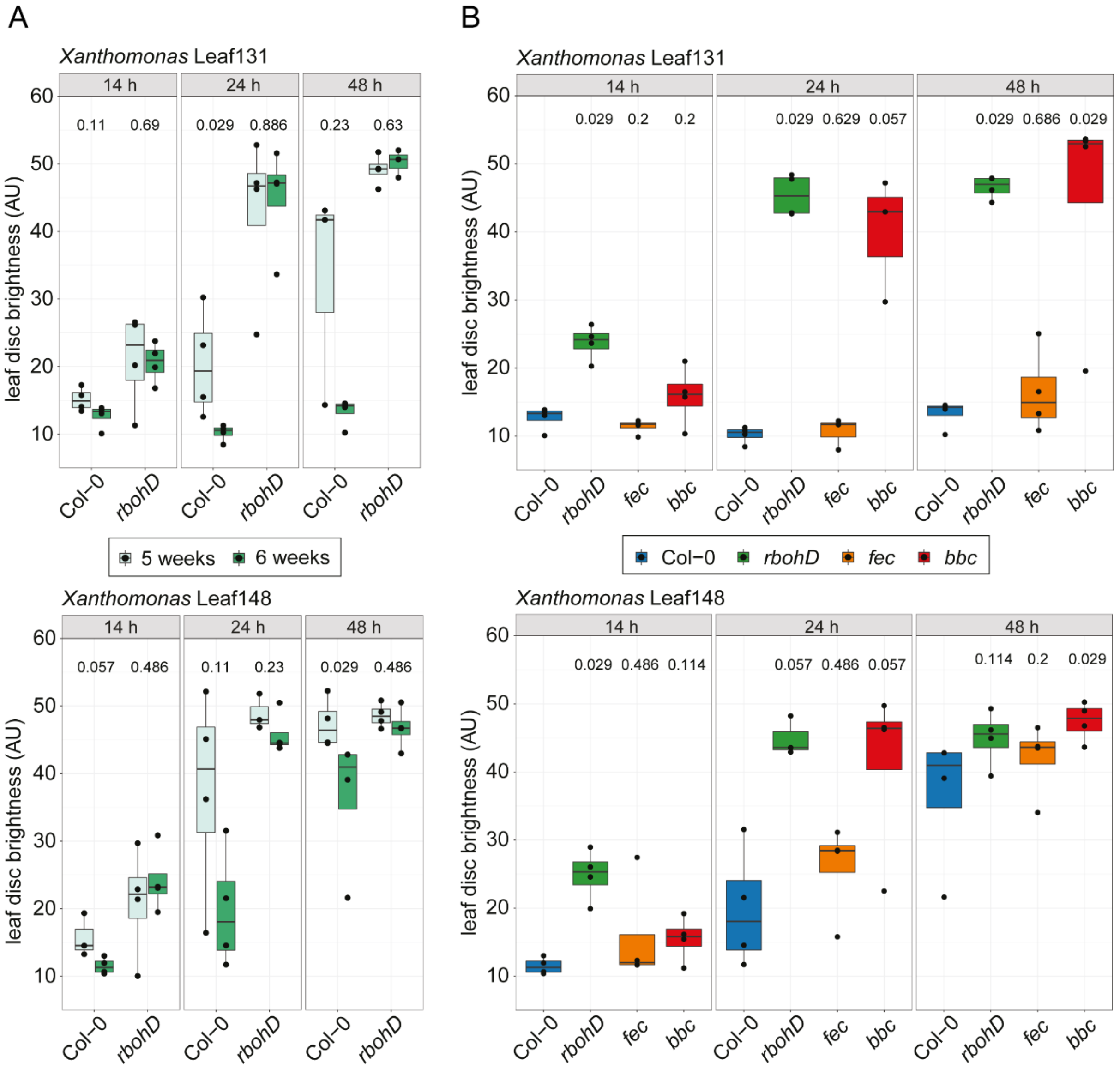


**Figure S3. Plant age and genotype influence leaf disc degradation by *Xanthomonas*. A)** Time-course of leaf discs brightness from five- and six-week-old Col-0 and *rbohD* plants inoculated with *Xanthomonas* Leaf131 (top panel) or Leaf148 (bottom panel). **B)** Time-course of leaf discs brightness from six-week-old Col-0, *rbohD, fls2wfrcwerk1 (fec)* and *bak1-5bkk1cerk1 (bbc)* plants inoculated with *Xanthomonas* Leaf131 (top panel) or Leaf148 (bottom panel). Statistical differences of leaf disc brightness between plant age (A) or plant genotype (B) at varying time points is indicated with p-value above box plots (two-tailed Mann–Whitney *U*-test, n = 4). Box plots show the median with upper and lower quartiles and whiskers present 1.5x interquartile range.

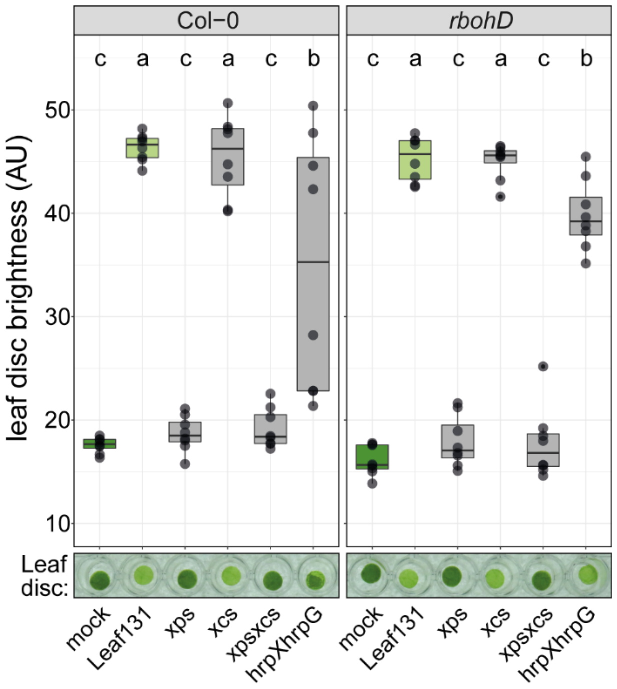


**Figure S4. *Xanthomonas* Leaf131 hrpX and hrpG are not necessary for leaf disc degradation.** Leaf discs of Col-0 or *rbohD* plants (five weeks old) were mock treated or with *Xanthomonas* Leaf131 wildtype or mutant strains and incubated for 24 hours. Significant differences were calculated with ANOVA and Tukey's HSD post hoc test (n= 8, letters indicate significance groups, $\alpha = 0.05$).

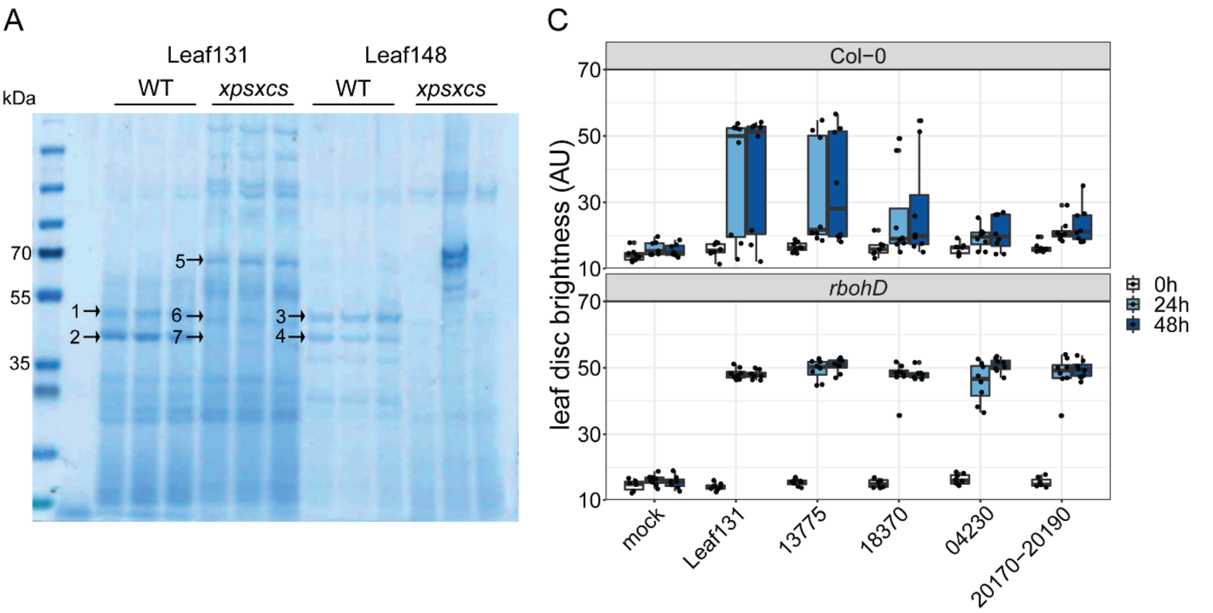


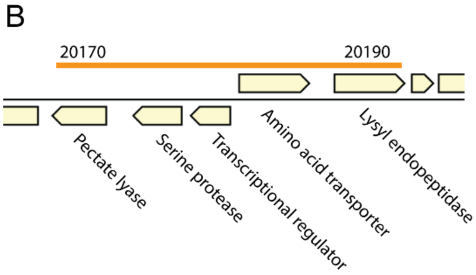


D

| Accession Number | Protein | Molecular Weight | Homologue of other *Xanthomonas* spp. |
|---|---|---|---|
| ASF73_13775 | Endoglucanase | 52 kDa | XCV0670[a], XOO4019[b] |
| ASF73_18370 | Serine protease | 64 kDa | XAC3980[c], XCV4074[d] |
| ASF73_04230 | Pectate lyase precursor | 40 kDa | - |
| ASF73_20170 | Pectate lyase precursor | 37 kDa | - |
| ASF73_20190 | Lysyl endopeptidase | 48 kDa | - |

[a] Sole et al. 2015, J. Bacteriol.
[b] Jha et al. 2007, MPMI
[c] Laia et al. 2009 BMC Microbiol.
[d] Szczesny et al. 2010, New Phytol.

**Figure S5. Proteomic analysis of supernatant from *Xanthomonas* Leaf131 and Leaf148 liquid culture identified T2SS-specific proteins. A)** Coomassie stained SDS-PAGE of cell-free supernatant from *Xanthomonas* Leaf131 and Leaf148 wildtypes and *xpsxcs* mutants liquid cultures (n=3). Black arrows with numbers indicate protein bands excised from gel and analysed by LC-MS/MS with results of differen fractions shown in Supplementary table 1. **B)** Genomic region encoding T2SS-dependent secreted proteins pectate lyase (ASF73_20170) and lysyl endopeptidase (ASF73_20190). Orange line indicates in-frame deletion of gene cluster. **C)** Leaf discs of Col-0 or *rbohD* plants (six weeks old) were mock treated or with *Xanthomonas* Leaf131 wildtype or mutant strains with gene deletions of ASF73_13775, ASF73_18370, ASF73_04230, ASF73_20170-ASF73_20190. **D)** Table shows candidate genes of T2SS-dependent secreted proteins identified by LC-MS/MS (Supplementary table 1) and gene identifiers of homologues in other *Xanthomonas* species as described in the literature.

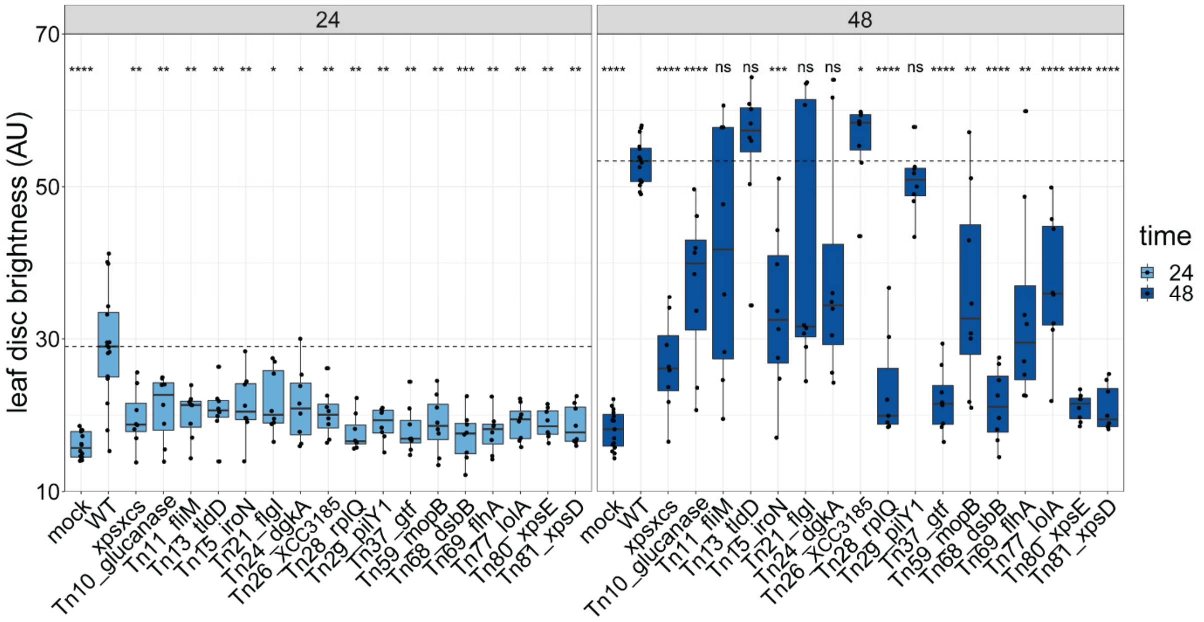


**Figure S6. Transposon mutagenesis screen revealed candidate genes involved in leaf degradation.** Leaf discs of five-week-old *rbohD* plants were mock inoculated (10 mM $MgCl_2$) or with *Xanthomonas* Leaf131 wildtype (WT) or Tn mutants (OD=0.02). Significant difference of Tn mutants compared to WT was determined by two-tailed Mann–Whitney *U*-test (n = 8) and p-values indicated as ns, non-significant; *, p<0.05, **, p<0.01; ***, p<0.001; ****, p<0.0001. Dashed line shows median of WT control.

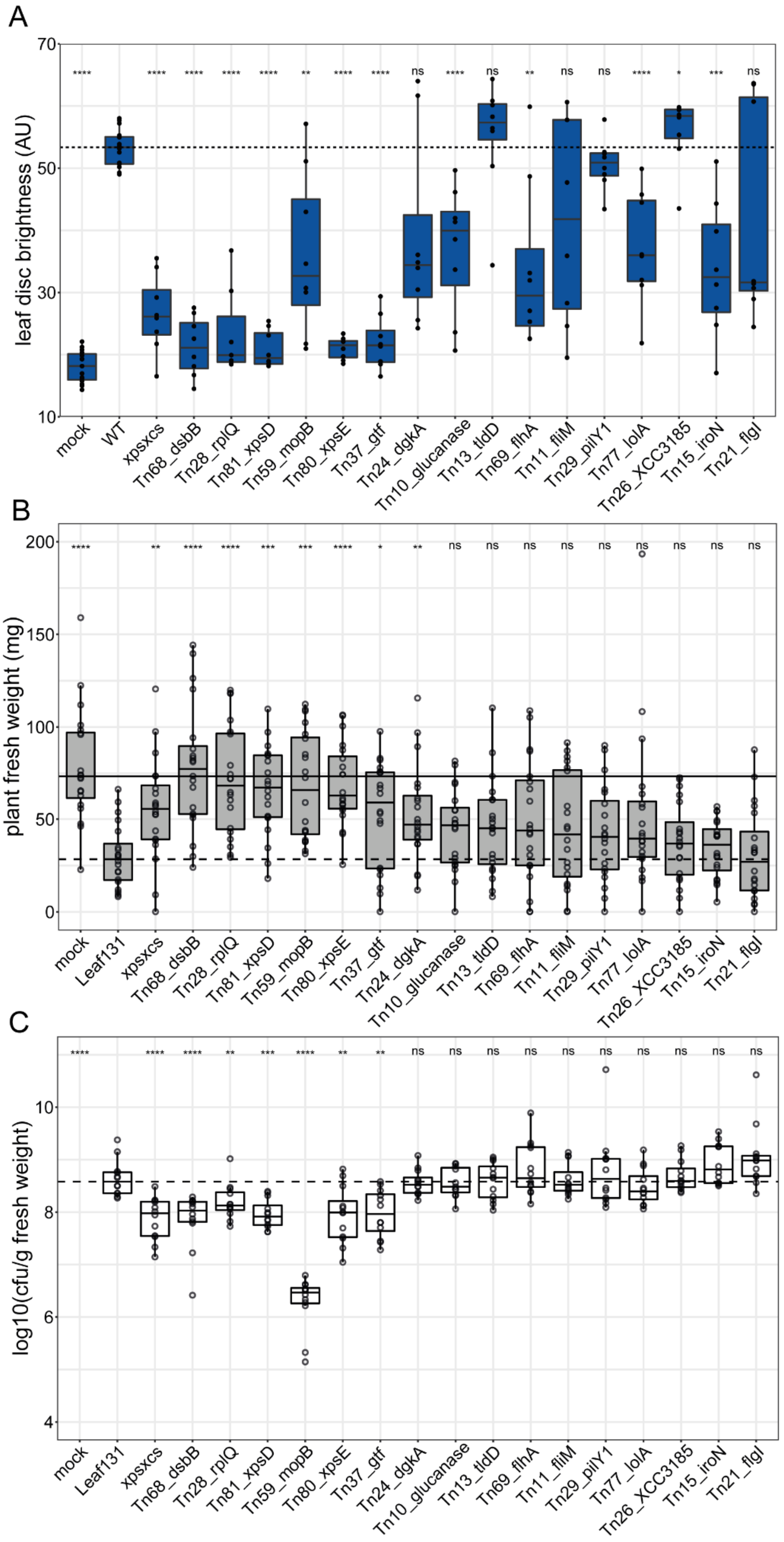


**Figure S7. Virulence of *Xanthomonas* Leaf131 transposon mutant candidates. A)** Leaf discs of five-week-old *rbohD* plants were mock inoculated (10 mM $MgCl_2$) or with *Xanthomonas* Leaf131 wildtype (WT) or Tn mutants (OD=0.02) and incubated for 48 hours (same data as shown in Figure

S5, 48 hours). Significant difference of Tn mutants compared to WT was determined by two-tailed Mann–Whitney *U*-test (n = 8) and p-values indicated as ns, non-significant; *, p<0.05, **, p<0.01; ***, p<0.001; ****, p<0.0001. **B)** Fresh weight of aboveground plant tissue of five-week-old gnotobiotic *rbohD* plants either mock inoculated or with *Xanthomonas* Leaf131 wildtype or Tn mutants. **C)** Colony forming unit (CFU) counts of *Xanthomonas* Leaf131 per gram plant fresh weight from samples in B). Box plots show the median with upper and lower quartiles and whiskers present 1.5× interquartile range. Significant differences in B) (n=15) and C) (n=12) were calculated by two-tailed Mann–Whitney *U*-test and p-values indicated as ns, non-significant; *, p<0.05, **, p<0.01; ***, p<0.001; ****, p<0.0001. Dashed line and solid line show median of WT control and mock control, respectively.

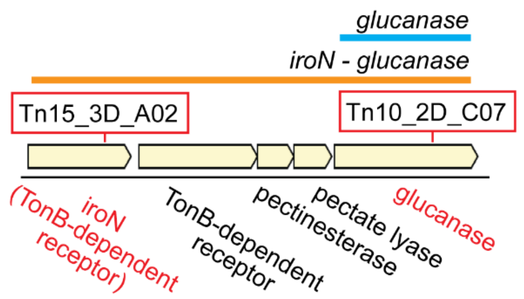


**Figure S8. Genomic region of *iroN* and *glucanase* in *Xanthomonas* Leaf131.** Red boxes indicate transposon insertion in indicated Tn mutants. Orange line highlights multi-gene deletion region in strain *iroN-glucanase*. Blue line indicates single gene deletion of *glucanase*.

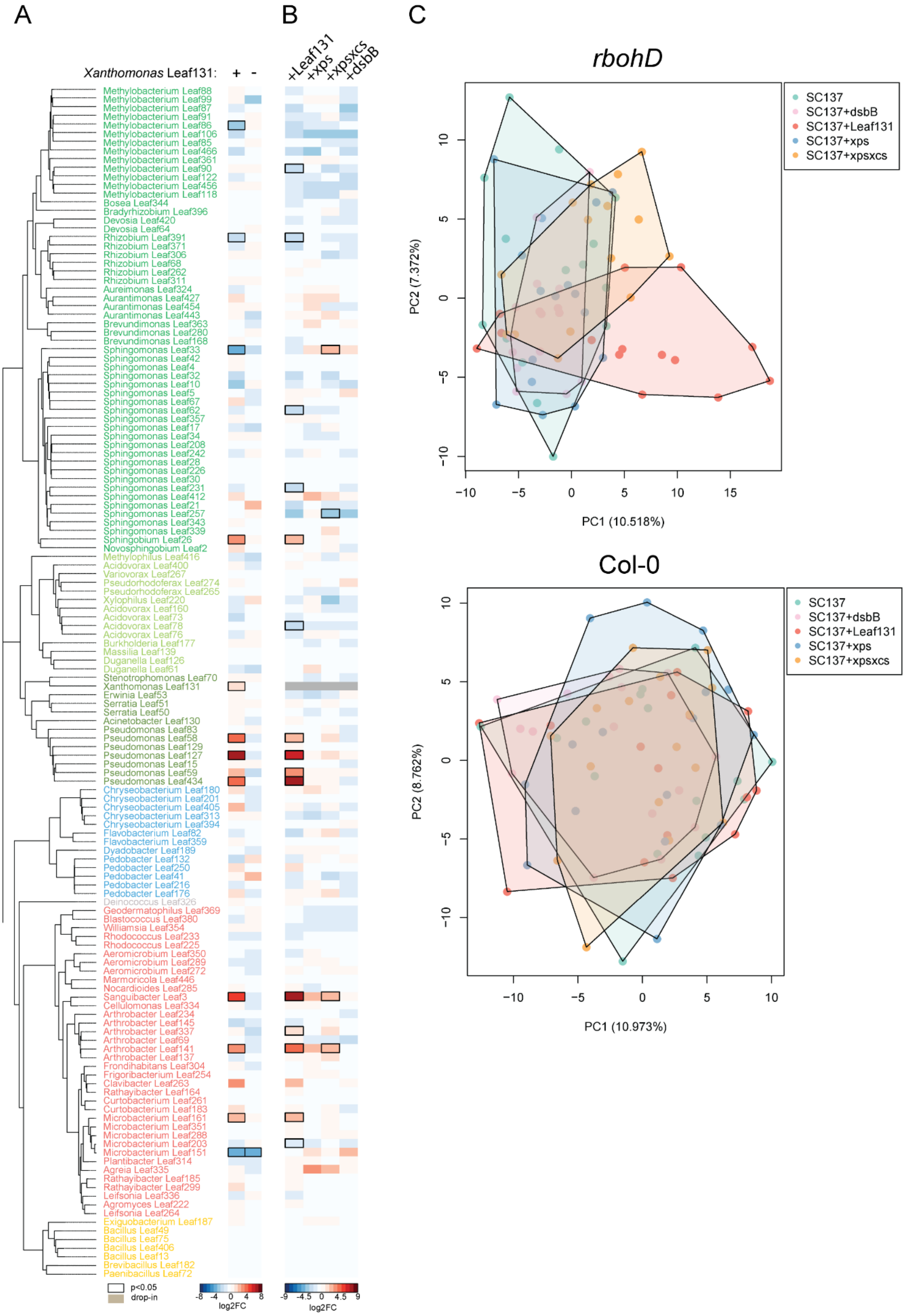


**Figure S9. Microbiota composition in presence of virulent or attenuated *Xanthomonas* Leaf131.** **A)** Heatmap shows $\log_2$ fold changes (log2FC) of strains in SynCom-137 in *rbohD* compared to Col-0

wild-type plants in the presence (+) or absence (-) of *Xanthomonas* Leaf131. **B)** Heatmap shows $\log_2$ fold changes (log2FC) of strains in the presence of either *Xanthomonas* Leaf131 wildtype or the mutants *xps*, *xpsxcs*, or *dsbB* compared to SynCom-137 without Leaf131. Black rectangles show significant changes, p-value < 0.05 (n = 16, Wald test, Benjamini–Hochberg adjusted). Subset of same data is shown in Figure 6A. **C)** Principal component analysis of community in *rbohD* plants (upper panel) and Col-0 (lower panel) inoculated only with SynCom-137 or SynCom-137 containing either *Xanthomonas* Leaf131, *xps*, *xpsxcs*, or *dsbB* in Axes show principal components PC1 and PC2 with their explained variance (%).

### Supplementary table legends

**Supplementary table 1. Proteomics of Leaf131 culture supernatant. A)** Identification of proteins in fractions of *Xanthomonas* Leaf131 supernatant from wildtype (fraction 1 and 2) or *xpsxcs* mutant (fraction 4 and 5). **B)** Identification of proteins in fractions of *Xanthomonas* Leaf148 supernatant from wildtype (fraction 3 and 4). Fractions are protein bands excised from SDS-PAGE (Figure S5A). Selection for gene knockout highlighted in orange. **C)** Table shows selected protein candidates for gene knockout in *Xanthomonas* Leaf131.

**Supplementary table 2. Transposon mutagenesis screen in *Xanthomonas* Leaf131. A)** Overview of transposon screen and identified candidate genes. **B)** Selection of validated candidate genes.

**Supplementary table 3. Knockout strains used in this study.**

**Supplementary table 4. Oligonucleotides used in this study.**

**Supplementary table 5. SynCom strains and microbiota composition data. A)** *At*-LSPHERE strains used in SynCom-137 and *Xanthomonas* Leaf131 and Leaf148. **B)** ASV count table for drop-out experiment and **C)** corresponding metadata. **D)** ASV count table for drop-in experiment and **E)** corresponding metadata.
